# Artificially stimulating retrotransposon activity increases mortality and accelerates a subset of aging phenotypes in *Drosophila*

**DOI:** 10.1101/2022.05.23.493120

**Authors:** Joyce Rigal, Ane Martin Anduaga, Elena Bitman, Emma Rivellese, Sebastian Kadener, Michael T. Marr

## Abstract

Transposable elements (TE) are mobile sequences of DNA that can become transcriptionally active as an animal ages. Whether TE activity is simply a byproduct of heterochromatin breakdown or can contribute towards the aging process is not known. Here we place the TE *gypsy* under the control of the UAS GAL4 system to model TE activation during aging. We find that increased TE activity shortens the lifespan of male *D. melanogaster.* The effect is only apparent in middle aged animals. The increase in mortality is not seen in young animals. An intact reverse transcriptase is necessary for the decrease in lifespan implicating a DNA mediated process in the effect. The decline in lifespan in the active *gypsy* flies is accompanied by the acceleration of a subset of aging phenotypes. TE activity increases sensitivity to oxidative stress and promotes a decline in circadian rhythmicity. The overexpression of the Forkhead-box O family (FOXO) stress response transcription factor can partially rescue the detrimental effects of increased TE activity on lifespan. Our results provide evidence that active TEs can behave as effectors in the aging process and suggest a potential novel role for dFOXO in its promotion of longevity in *D. melanogaster*.

## INTRODUCTION

Aging leads to a progressive loss of physiological integrity that culminates in a decline of function and an increased risk of death (López-Otín et al., 2013). It is a universal process that involves the multifactorial interaction of diverse mechanisms that are not yet fully elucidated. Transposable elements (TE) are among the many factors that have been proposed to be involved in aging (Morley, 1995; Wood & Helfand, 2013). TE are present in every eukaryotic genome sequenced to date (García Guerreiro, 2012; Huang et al., 2012). They are sequences of DNA that can move from one place to another (McCLINTOCK, 1950), either by reverse transcription and insertion into the genome (Class 1: Retrotransposons), or through direct excision and movement of the element (Class 2: DNA TE) (McCullers & Steiniger, 2017).

Multiple studies (Gorbunova et al., 2021) report that TE mRNA levels increase in the aging somatic tissue of flies (Giordani et al., 2021; Li et al., 2013; Wood et al., 2016), termites (Elsner et al., 2018), mice (De Cecco, Criscione, Peterson, et al., 2013), rats (Mumford et al., 2019) and humans (LaRocca et al., 2020). A direct correlation to an increase in genomic copy number has been difficult to determine (Treiber & Waddell, 2017; Yang et al., 2022). Nonetheless, clear evidence for an increase of TE somatic insertions with age has been obtained using reporter systems. Two reporter systems for insertions of the long-terminal-repeat (LTR) retrotransposon *gypsy* demonstrate that *gypsy* insertions increase during aging in the *D. melanogaster* brain and fat body (Y.-H. Chang et al., 2019; Li et al., 2013; Wood et al., 2016).

TE movement in somatic tissue has been proposed to be a driver of genomic instability (Ivics & Izsvák, 2010) and potentially aging (Morley, 1995; Wood & Helfand, 2013; Woodruff & Nikitin, 1995). Additionally, TE activity has been reported to cause disease in humans (Hancks et al., 2012). Current research reports that long-interspersed-element-1 (*L1)* activity itself, without an increase in insertions, can trigger an inflammation response that contributes to aging related phenotypes in human senescent cells (Cecco et al., 2019). In aged mice, the use of reverse transcriptase inhibitors can downregulate this age associated inflammation (Cecco et al., 2019) implicating retrotransposons. The shortened lifespan of a Dicer-2 (*dcr-2)* mutant fly strain, which has an increase of TE expression, can also be extended by the use of reverse transcriptase inhibitors (Wood et al., 2016). In summary, current evidence suggests a role for TE activity in aging.

However, whether the role of TE activity is as effector or bystander of the aging process is an open question. As an animal ages, heterochromatin repressive marks decrease and resident silent genes can become expressed (Jiang et al., 2013). TE sequences are enriched in silent heterochromatin and thus become expressed (De Cecco, Criscione, Peckham, et al., 2013). This raises the question of whether TE expression is simply a byproduct of age-related heterochromatin breakdown or if TE themselves can contribute to the aging process. To date, this has not been directly assayed.

To combat the effects of TE activity, cells have evolved small RNA pathways to maintain silencing of TE. The PIWI pathway dominates in the germline while the somatic tissue of *Drosophila* is thought to mainly rely on the siRNA pathway (Hyun, 2017). This pathway is based on Dicer-2 cleaving double stranded (ds) RNA precursors, generally viral genomes or TE dsRNA, into small RNAs that are loaded into Ago2 guiding the RNA Induced Silencing Complex (RISC) to cleave its targets (Hyun, 2017). In *D. melanogaster*, endogenous siRNAs in RISC mapping to TE loci have been reported (Czech et al., 2008; Ghildiyal et al., 2008; Kawamura et al., 2008). The mutation, knockdown, or depletion of genes involved in the siRNA pathway such as *Dcr*-2 (Lee et al., 2004) and *AGO2* (Okamura et al., 2004) leads to increased expression of TE in somatic cells and a shortened lifespan (Kawamura et al., 2008; Czech et al., 2008; Chung et al., 2008; Lim et al., 2011; Li et al., 2013; Wood et al., 2016; Chen et al., 2016). On the other hand, the overexpression of *Dcr-2* (Wood et al., 2016) and *AGO2* (Yang et al., 2022) in adult fly somatic tissue can lower TE expression and extend lifespan. Taken together, these results suggest TE activity can influence lifespan. The effect of TE activity on lifespan has not been directly determined.

The Forkhead-box O family (FOXO) is an evolutionarily conserved transcription factor capable of enhancing longevity by enabling the cell to respond to diverse stress signals (Calnan & Brunet, 2008). The longevity effects of FOXO have been reported in worms, flies, and mice (Martins et al., 2016). FOXO activity can enable transcriptional responses to provide protective effects against cellular stress: oxidative stress, heat shock, virus infection, and defects in protein homeostasis among many others (Donovan & Marr, 2016; Martins et al., 2016; Spellberg & Marr, 2015). Whether TE activity is a condition that FOXO activity might protect against is unknown.

In this study we have set up a system to directly assay if TE can become active contributors of the aging process. For this, we placed the retrotransposon *gypsy* (Bayev et al., 1984), which will be used as a candidate to model TE activity during aging, under the control of the UAS GAL4 system (Brand & Perrimon, 1993). The retrotransposon *gypsy* is a TE that has been clearly shown to become active in aged *D. melanogaster* somatic tissue under natural conditions (Li et al., 2013; Wood et al., 2016). In our system, we find that active TE expression significantly increases the mortality in middle aged flies and that an intact reverse transcriptase is necessary for this effect. The increase in mortality is accompanied by acceleration of a subset of aging-related phenotypes. We find that the FOXO homologue in *Drosophila* (dFOXO) can counteract and partially rescue the decrease in lifespan generated by an active TE, suggesting a possible novel mechanism through which dFOXO might promote longevity in *D. melanogaster*.

## RESULTS

To determine dFOXO’s possible involvement in TE regulation we sequenced the RNA from whole animals of 5-6 days (d) and 30-31d old *w^DAH^*(wt) and dFOXO deletion (*w^DAH^ ^Δ94^*) males (Slack et al., 2011). These lines have been extensively backcrossed, making them isogenic other than for the dFOXO deletion (Slack et al., 2011). In these conditions, the wt animals display significant increased expression of two TEs (*1731* and *Doc4*) and decreased expression of two additional TEs (*Stalker2* and *G6*) with age (Fig. 1a, 1b and 1-S1). By contrast, in the dFOXO deletion animals, 18 TEs exhibited a significant increase in expression with age (Fig. 1c, 1d and 1-S2) while no TE expression decreased with age indicating the overall TE load is greater in the aged dFOXO deletion flies.

**Figure 1.**
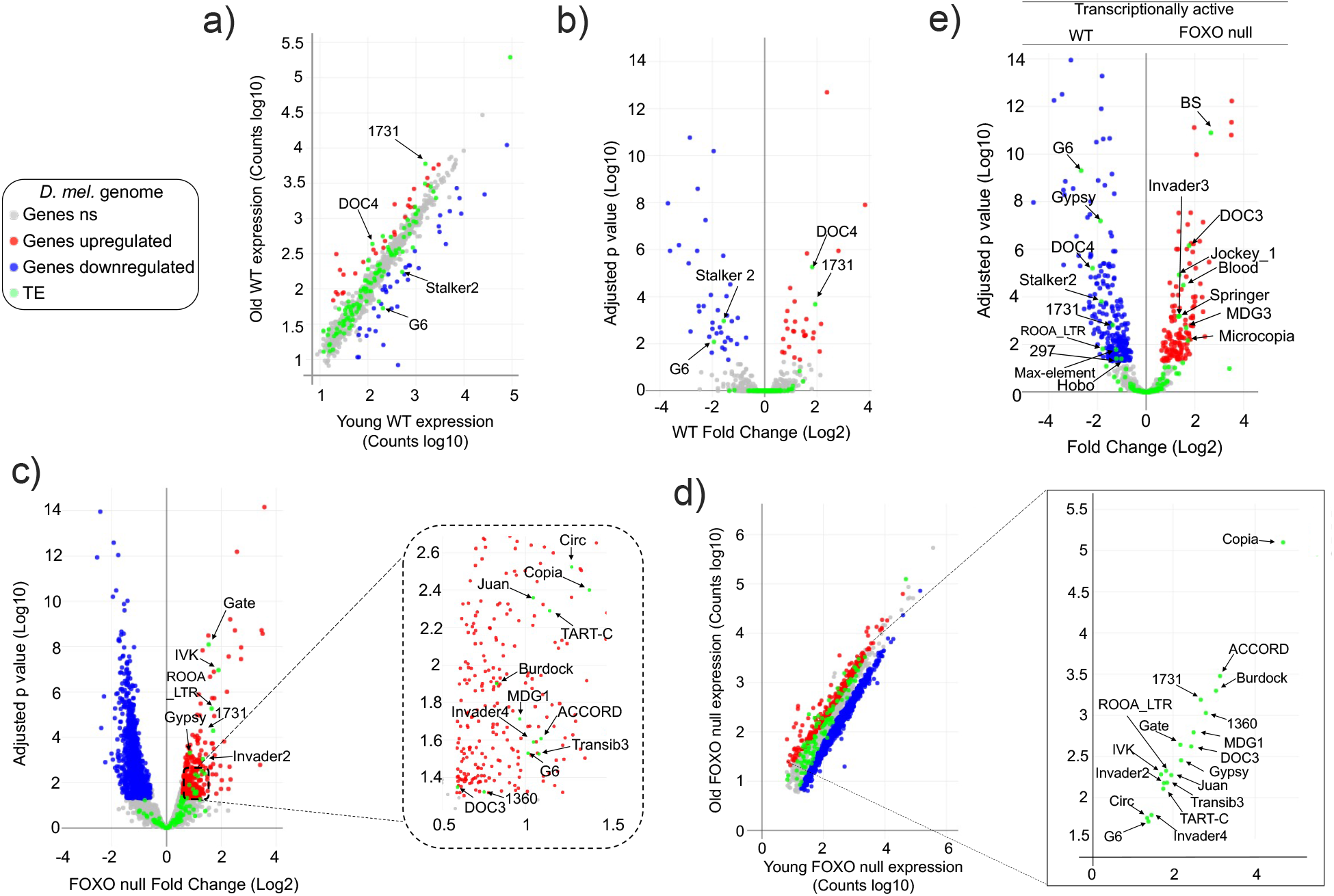
TE expression increases with age in FOXO null flies. Data represents RNAseq from 3 biological replicates (10 male flies per sample). Young – 5 days, Old – 30 days. Legend (grey: fly genes not significant (ns), red: upregulated fly genes, blue: downregulated fly genes, green: TE). Differential expression indicates a 1.5-fold change or higher and an adjusted p value < 0.05, as determined by DEseq2. a-d) Red indicates upregulation with age. Blue denotes downregulation with age. Significantly different TE are identified by name. a-b) Wildtype (WT) control. 4 TE are labelled. c-d) FOXO null. 18 TE are labelled. e) Volcano plot of young WT control and young FOXO null. Red and blue dots indicate gene relative expression in FOXO null compared to WT, respectively. TE are marked by green dots. Adjusted p values for each TE are provided in Figure 1-Table supplement 1 (WT) and Figure 1-Table supplement 2 (FOXO null)

The vast majority of TE expression levels in both strains fall within the observed range of average gene expression (Fig. 1a and Fig. 1d). Total TE expression undergoes a small change overall in both genotypes. Expression increased only 1.2-fold with age (Fig. 1-Table S1) in wt flies and 1.41-fold in dFOXO null flies (Fig. 1-Table S2). Of the two TE that increased with age in wt flies, only *1731* exhibits an increase in dFOXO null flies. Among the two TE that decreased with age in wt, *G6* showed the opposite effect on expression in dFOXO null flies while *Stalker2* showed no change with age in dFOXO null flies. The direct comparison of individual TE expression levels in young wt and dFOXO null flies indicates that despite being backcrossed (Slack et al., 2011) different TE are being expressed in each strain (Fig. 1e). This means that the otherwise isogenic lines have a different transcriptionally active TE landscape. Therefore, beyond the comparison of the number of transcriptionally active TE, a direct comparison between wt and dFOXO null flies to determine the effect of dFOXO on any specific TE expression during aging is challenging.

Our difficulties to determine the effect of dFOXO on any specific TE highlights the need for a controlled system to test TE expression and regulation during aging. The *gypsy* retrotransposon was selected as a model system to study TE activity and we developed a UAS-*gypsy* system. The structure of the ectopic UAS-*gypsy* system is depicted in Fig. 2a. A direct comparison to the endogenous *gypsy* element can be observed in a simplified overview of three broad stages in the *gypsy* lifecycle (parental element, propagation through transcription and reverse transcription, and new insertion). The presence of the UAS promoter in the ectopic *gypsy* allows the control of *gypsy* expression by mating to a GAL4 expressing strain. In addition, we inserted a unique sequence tag in the 3’ LTR of the ectopic *gypsy* to differentiate it from the endogenous copy of *gypsy* (Bayev et al., 1984, p. 4). We inserted the UAS-*gypsy* construct in the VK37 attP site on chromosome 2 using the PhiC31 system (Venken et al., 2006). Using this approach, we found that ectopic *gypsy* expression is significantly induced when the line is crossed to a strain in which gal4 is ubiquitously expressed under the *ubiquitin* promoter (*Ubi*>*gypsy*) (Fig. 2b).

**Figure 2.**
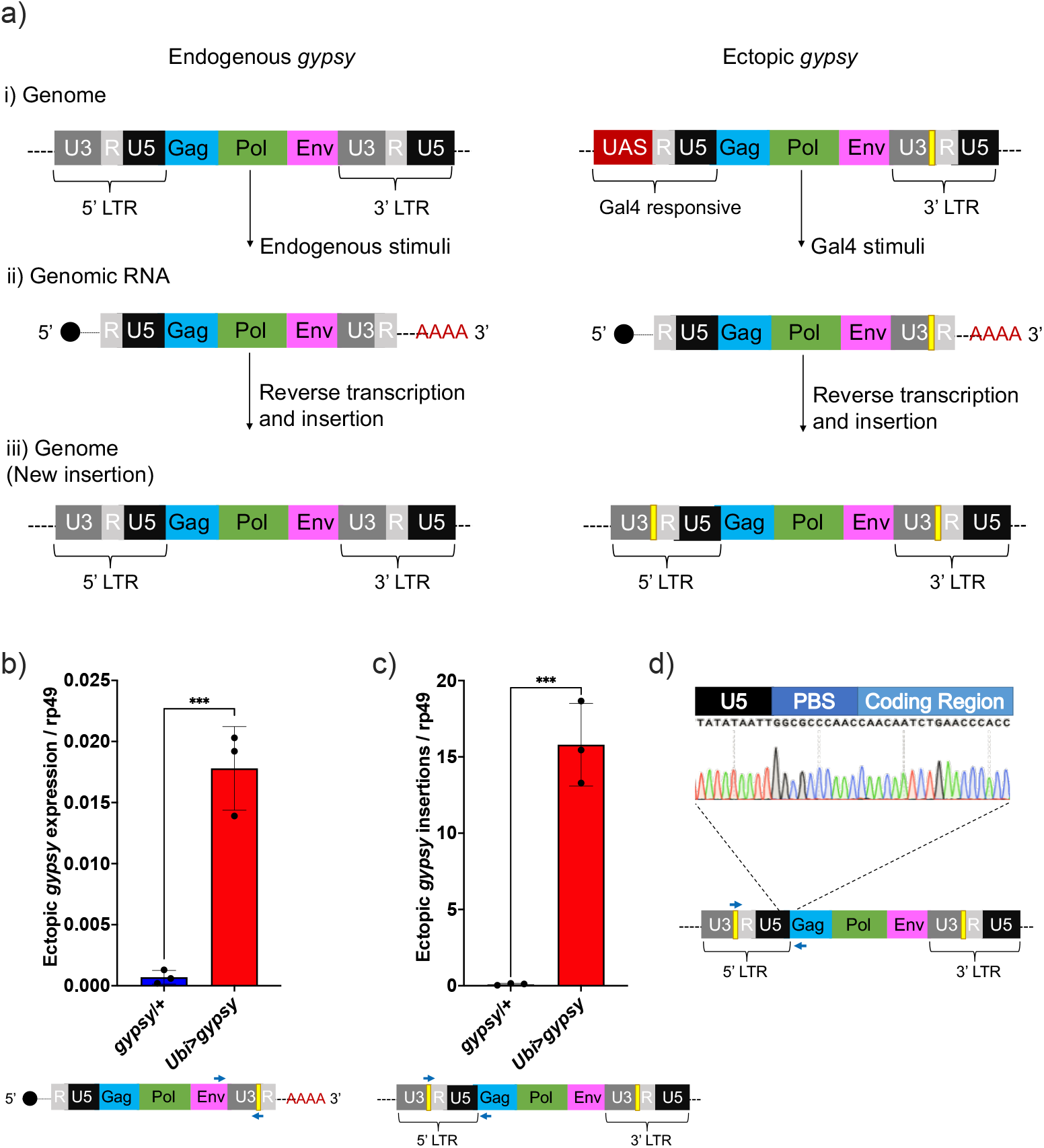
UAS-*gypsy* structure and functional test. a) Simplified overview of 3 stages of the *gypsy* retrotransposon lifecycle, i) parent insertion in the genome, ii) the transcribed RNA (genomic RNA), and iii) the new copy inserted in the genome. The difference between the wildtype *gypsy* and ectopic *gypsy* resides in both its 5’ and 3’ LTR. The presence of an upstream activating sequence (UAS) in the 5’ LTR allows *gypsy* to be transcribed in response to a Gal4 stimuli. In the 3’ LTR, the addition of a unique sequence of DNA (denoted by a yellow square) not found in the *D. melanogaster* genome allows quantification and tracking of new insertions by molecular methods. b) RT-qPCR of 5-day old males. 3’ end of ectopic *gypsy* transcript is detected. Data are represented as means ± SD (3 biological replicates, each dot is a pool of 5 flies). One-tailed t test, *** p value 0.0005. c) gDNA qPCR of 5-day old males. Ectopic *gypsy* provirus insertion junctions are detected. Data are represented as means ± SD (3 biological replicates, each dot is a pool of 10 flies). One-tailed t test, *** p value 0.0003. d) Sanger sequencing of the newly created ectopic *gypsy* provirus insertion junction in *Ubi*>*gypsy* flies.

This tag also allows us to take advantage of the strand transfer that occurs during reverse transcription of the retroelement before integrating into a new site in the genome. This process replaces the UAS sequence with a 5’ LTR and transfers the sequence tag to the new LTR of the newly integrated *gypsy* element (Weaver, 2008) (Fig. 2a). Using this approach, we found that new insertions are created and can be specifically detected (Fig. 2c) when UAS-*gypsy* is expressed (*Ubi*>*gypsy* genotype). The junction between the recombinant *gypsy* 5’ LTR and the *gypsy* provirus sequence is only formed when a complete insertion is made. Sanger sequencing of the qPCR insertion products demonstrates the specific detection of this junction, indicating that the UAS-*gypsy* element can transpose and is an active TE element (Fig. 2d).

To characterize the spectrum of insertion sites of the UAS-*gypsy* construct we took a targeted sequencing approach. Using a biotinylated primer hybridizing with the unique sequence tag oriented to read out from the newly integrated 5’ LTR we created Illumina Next generation sequencing libraries to sequence the genomic junctions (Fig. 3-S1a). Libraries were prepared from three biological replicates of ten 14-day old male flies each. Sequencing reads were sorted using the LTR sequence as a barcode. After removing the LTR sequence, the exact site of insertion was mapped back to the *Drosophila* reference genome. More than 11,000 insertion sites were identified. Sites were identified in all chromosomes with the fraction of insertions roughly correlating with the size of the chromosome (Fig. 3A). Comparing the insertion sites with the genome annotation allowed us to classify the insertion sites. Sixty-six percent of the mapped insertion sites are in transcribed regions (Fig. 3B). Of those sites, the majority are in intronic regions (Fig. 3C). Because the data provides nucleotide resolution, we could also determine the six-nucleotide target site duplication that occurs at the junction between the *gypsy* LTR and the genome upon insertion (Dej et al., 1998). The target site duplication consensus determined from our mapped insertion sites matches the known YRYRYR previously identified for *gypsy* integration sites (Figure 3D) (Dej et al., 1998). To determine the distribution of the insertions sites we divided the largest chromosome arms (3R, 3L, 2R, 2L, X) into roughly 1 megabase bins and counted the number of insertions per bin. Then we plotted the fraction of the total insertions on that chromosome arm in each bin. A plot for the arms of chromosome 3 is shown in figure 3E. The plots of the other chromosomes are in the supplemental figure 3-S1b. At this level of resolution, insertions are detected roughly evenly across the chromosomes.

**Figure 3.**
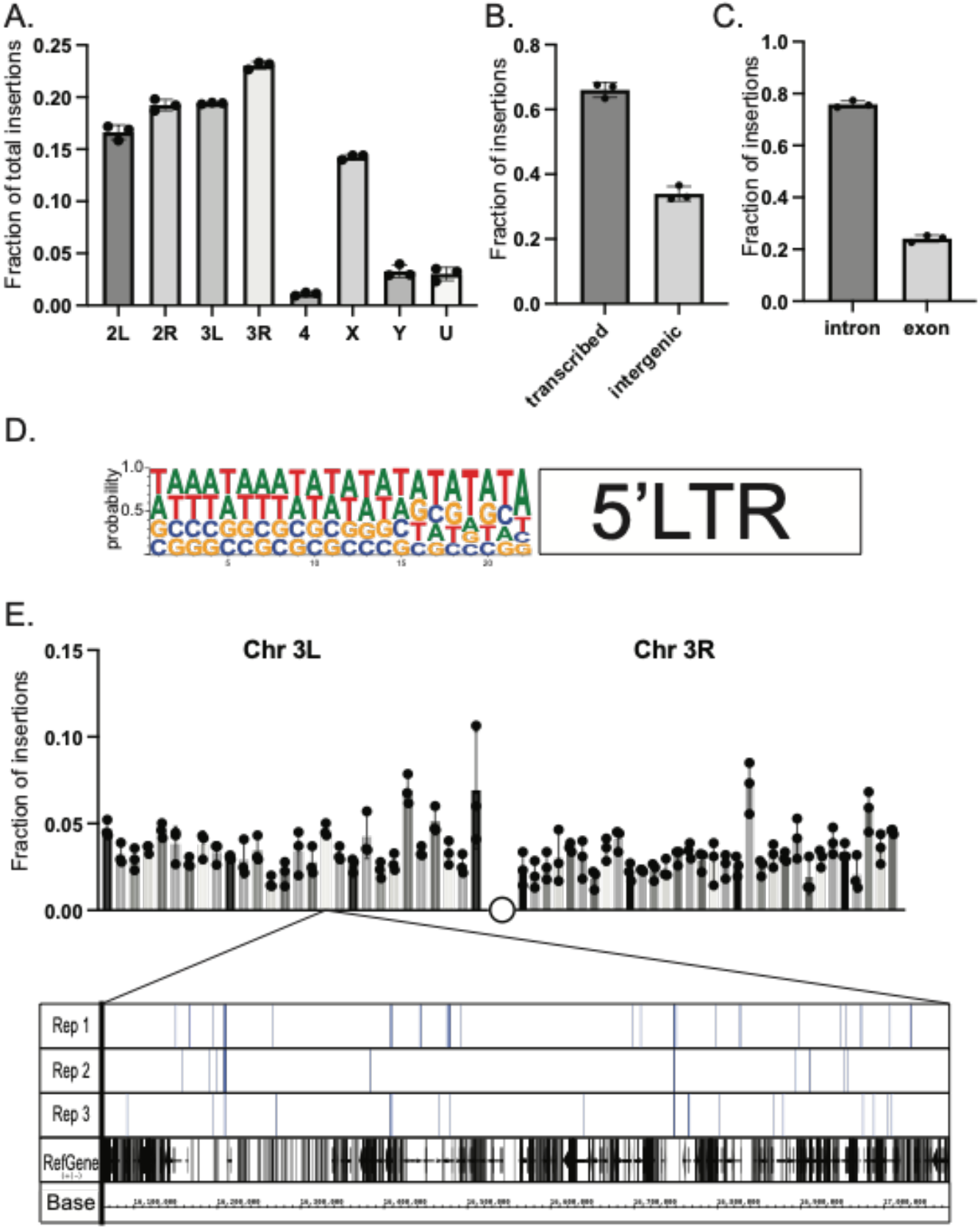
NGS mapping of ectopic *gypsy* insertions. **A.** The average fraction of total insertions is shown for each chromosome (4, X, Y) or chromosome arm (2L, 2R, 3L, 3R). In addition, the fraction mapping to unplaced contigs is indicated as U. **B.** The fraction of total reads mapping to transcribed regions of the genome and intergenic regions is graphed. **C.** The fraction of insertions that map to the transcribed regions of the genome are subdivided. Insertions mapping to regions annotated as introns or exons is graphed. **D.** The sequences of the junction of the new 5’ LTR and the *Drosophila* genome were aligned and used to determine the probability of finding each base at each position. These probabilities are indicated by the size of the letter at each position. **E.** The fraction of insertions that map to each one megabase region of the reference genome for the arms of chromosome 3 are plotted. For illustration, a genome browser view of the 1Mb in Chr3L is shown. The insertion sites in that region for each replicate is indicated. A collapsed track showing genes in that region is also shown. For all histograms, the bars represent the average of three biological replicates. Error bars indicate the standard deviation and the filled circles indicate the individual measurements. Each replicate is a pool of 10 male 14 day-old flies

The feasibility of the system allowed us to test whether TE activity could affect lifespan. We set up three independent cohorts to measure lifespan. The assays were performed at different times of the year. We consistently found that the somatic expression of an active *gypsy* significantly decreased the lifespan of male flies (Fig. 4a). The individual cohorts showed a consistent effect on lifespan and the merged data was used to analyze the effect. A 19% reduction of lifespan in the active *gypsy* male flies is observed compared to parental controls, with a median survival of 70 days and 86 days, respectively. A lifespan effect was also observed in females, surprisingly it was also present in the UAS-*gypsy* parental control (Fig. 4-S1). Interestingly, a molecular assay demonstrates that more detected insertions are present in female UAS-*gypsy* controls than their male equivalents (Fig. 4-S2). An additional molecular assay was done in the *Ubi*>*gypsy* genotype to look at the distribution of detected insertions between the head and body (Fig. 4-S3). No significant difference was detected in male flies, indicating an equivalent level of insertions throughout somatic tissue. In females, however, a significant bias toward insertions in the body was detected, indicating possible differential expression of UAS-*gypsy* in the mixed tissues. Due to the confounding effects of the system observed in female flies (lifespan defect in parental UAS-*gypsy* control and differential distribution of detected insertions in *Ubi*>*gypsy*) male flies were used for all following experiments. The lifespan effect on the *Ubi*>*gypsy* flies only becomes apparent once the flies start to age. At 26 days old the survival curves noticeably separate from the parental controls and calculation of the age specific mortality shows a significant increase in mortality present in 26 to 75 day old animals (Fig. 4b-e).

**Figure 4.**
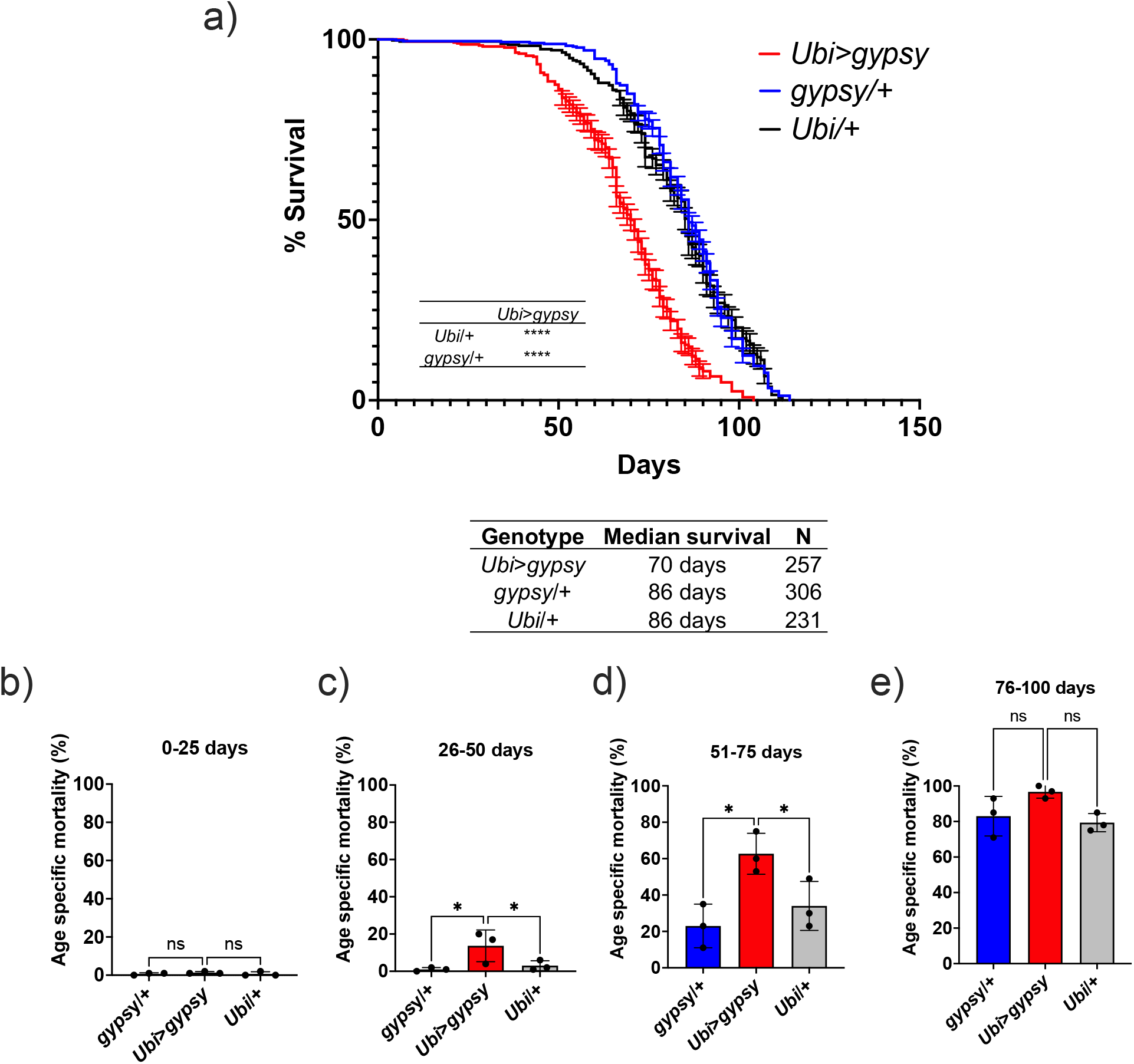
Ectopic *gypsy* expression decreases lifespan during old age. a) Survival curves of male flies expressing *gypsy* under the control of *Ubiquitin* Gal4 (Red) and the parental controls: *gypsy*/+ (Blue) and *Ubi*/+ (Black). Data represents 3 biological replicates (independent cohorts done at different times of year), error bars SE. **** p value <0.0001, Log-rank test. b-e) Data are represented as means ± SD (individual measurements are shown as dots, age specific mortality was calculated for each cohort independently). One-way ANOVA, adjusted p value: 26-50 days (*gypsy*/+ 0.047, *Ubi*/+ 0.047), adjusted p value 51-75 days (*gypsy*/+ 0.015, *Ubi*/+ 0.030), ns (not significant).

Ectopic *gypsy* DNA was detected at different ages in the survival curve (5, 14, 30, 50, and 70 days old) with three different primer pairs. In Figure 5a, the recombinant 5’ LTR *gypsy* junction is targeted to detect complete ectopic *gypsy* elements. In Figure 5b, the 3’ fragment of ectopic *gypsy* is targeted for detection. The 3’ fragment comprises the region between the *env* gene and the tag in the 3’ LTR. Complete and incomplete *gypsy* elements are quantified by this approach. In Figure 5c, the wildtype *gypsy* env region is targeted. This approach captures both the endogenous and ectopic *gypsy* content in the genome. Of the three approaches, the detection of the 5’ LTR *gypsy* junction (Fig. 5a) is the most stringent because only complete *gypsy* elements are detected. This is reflected in the smaller content of ectopic *gypsy* DNA detected compared to the 3’ fragment assay which detects both complete and incomplete *gypsy* elements (Fig 5b-c). The three approaches detect a relatively constant level of DNA while the animals are young with a surprising decrease in detection of ectopic *gypsy* DNA in older animals. Interestingly, ectopic *gypsy* RNA expression remains constant throughout the assayed timepoints (Fig. 5d). Although variability increases greatly with age.

**Figure 5.**
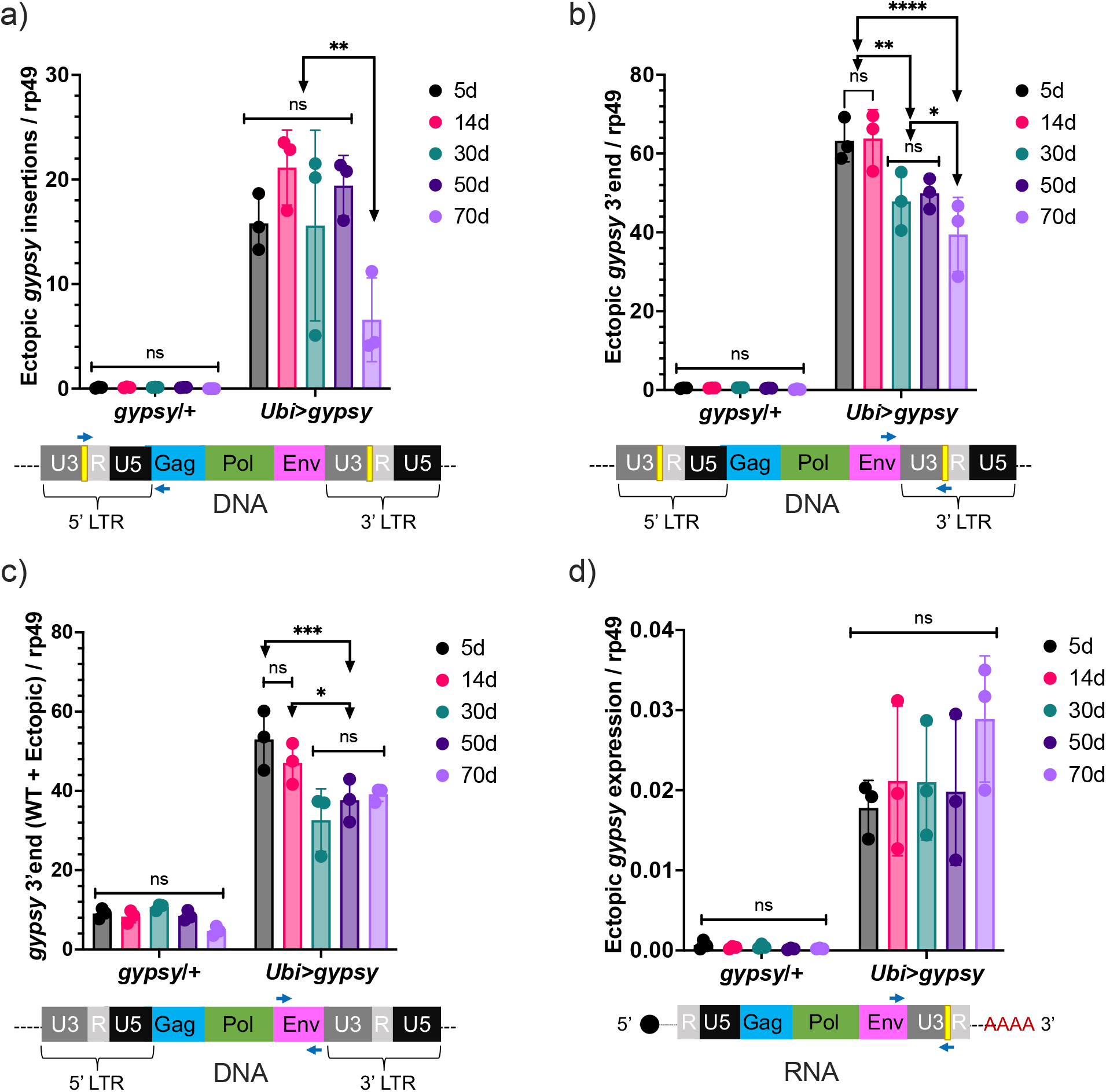
Ectopic *gypsy* DNA does not increase with age. a-c) gDNA qPCR of male flies at different ages (5d, 14d, 30d, 50d and 70d). Data are represented as means ± SD (3 biological replicates, each dot is a pool of 10 flies). 2way ANOVA, **** adjusted p value <0.0001, ** adjusted p value <0.01, * adjusted p value < 0.05. a) Ectopic *gypsy* provirus junctions are detected. b) 3’ end of ectopic *gypsy* fragments are detected. c) Wildtype (WT) and ectopic *gypsy env* fragments are detected. d) RT-qPCR of male flies at different ages (5d, 14d, 30d, 50d and 70d). 3’ end of ectopic *gypsy* transcript is detected. Data are represented as means ± SD (3 biological replicates, each dot is a pool of 5 flies). 2way ANOVA, not significant (ns).

The effect on lifespan caused by TE activity could be due to the process of retrotransposition through a DNA intermediate or by disruption of RNA homeostasis. One way to address this is to remove the DNA synthesis step and test if TE RNA presence alone can mediate the decrease in lifespan. This led us to test whether the lifespan effect would be present after deleting the reverse transcriptase (RT) from the UAS-*gypsy* polyprotein. We created a UAS-*gypsy* construct that contains an in-frame deletion in the polyprotein that removes most of the RT (Marlor et al., 1986) and inserted it in the same attP landing site on chromosome 2 (UAS-*ΔRT*). The deletion of RT in the UAS-*gypsy* transgene prevents the lifespan effect observed when crossed to *Ubi*-gal4 (*Ubi>ΔRT* genotype) (Fig 6a). This is not due to a defect in mRNA levels because RT-qPCR indicates that *Ubi>ΔRT* and *Ubi*>*gypsy* express the transposon RNA at the same level (Fig. 6b). Consistent with the loss of RT activity, we detect no increase in ectopic *gypsy* 3’ DNA in the *Ubi>ΔRT* line despite expressing equivalent levels of mRNA as *Ubi*>*gypsy* (Fig. 6c). This result indicates that an active RT is required for the observed decrease in lifespan seen when *gypsy* is ectopically expressed in somatic tissue.

**Figure 6.**
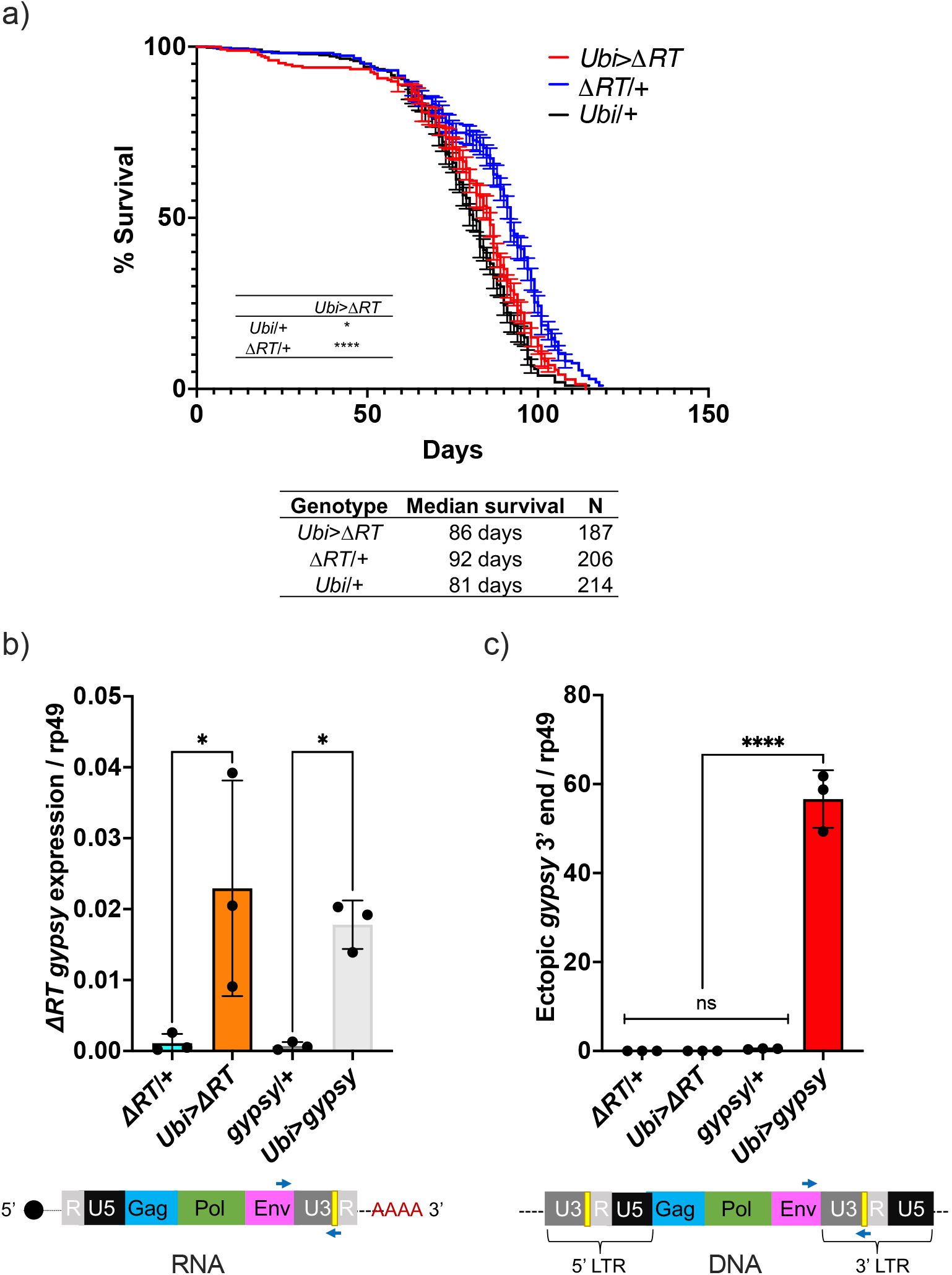
Decrease in lifespan requires reverse transcriptase activity. a) Survival curves of male flies expressing *ΔRT gypsy* under the control of *Ubiquitin* Gal4 (Red) and the parental controls: *ΔRT gypsy*/+ (Blue) and *Ubi*/+ (Black). Data represents 2 biological replicates (staggered independent cohorts), error bars SE. **** p value <0.0001, * p value 0.016, Log-rank test. b-c) Data are represented as means ± SD (3 biological replicates, each dot is a pool of 5 flies). *gypsy*/+ and *Ubi*>*gypsy* data are replotted from Fig. 2 b-c, respectively. b) RT-qPCR of 5-day old males. 3’ end of ectopic *gypsy* transcript is detected. One-way Anova, *ΔRT*/+ vs. *Ubi>ΔRT* * adjusted p value 0.012, *gypsy*/+ vs. *Ubi*>*gypsy* * adjusted p value 0.029. c) gDNA qPCR of 5-day old males. 3’ end of ectopic *gypsy* fragments is detected. Ordinary one-way Anova, **** adjusted p value <0.0001.

To determine if the decrease in lifespan of the active *gypsy* flies was accompanied by an acceleration of aging associated phenotypes, we decided to test if the ectopic expression of the *gypsy* TE resulted in the early emergence of any aging hallmarks. In particular, we focused on four different phenotypes that appear in normal aging flies (decreased resistance to dietary paraquat (Arking et al., 1991), decline in negative geotaxis (Rhodenizer et al., 2008), decrease of total activity (Sun et al., 2013) and disruption of circadian phenotypes (Curran et al., 2019; Rakshit et al., 2012).

For the paraquat resistance assays, 5, 15, 30 and 50 day old flies were exposed to 20 mM paraquat. As expected, resistance to paraquat exposure decreased with age in all genotypes (Fig. 7a). Interestingly, the survival curve of the 5-day old active *gypsy* flies mimicked that of the 15-day old parental control curves. Meanwhile the 15-day old active *gypsy* flies had a significant decrease in resistance to paraquat exposure when compared to age matched parental controls. However, once the flies are 30 and 50 days old it appears TE activity can no longer exacerbate the oxidative stress survival as experimental and control flies die at a similar rate.

**Figure 7.**
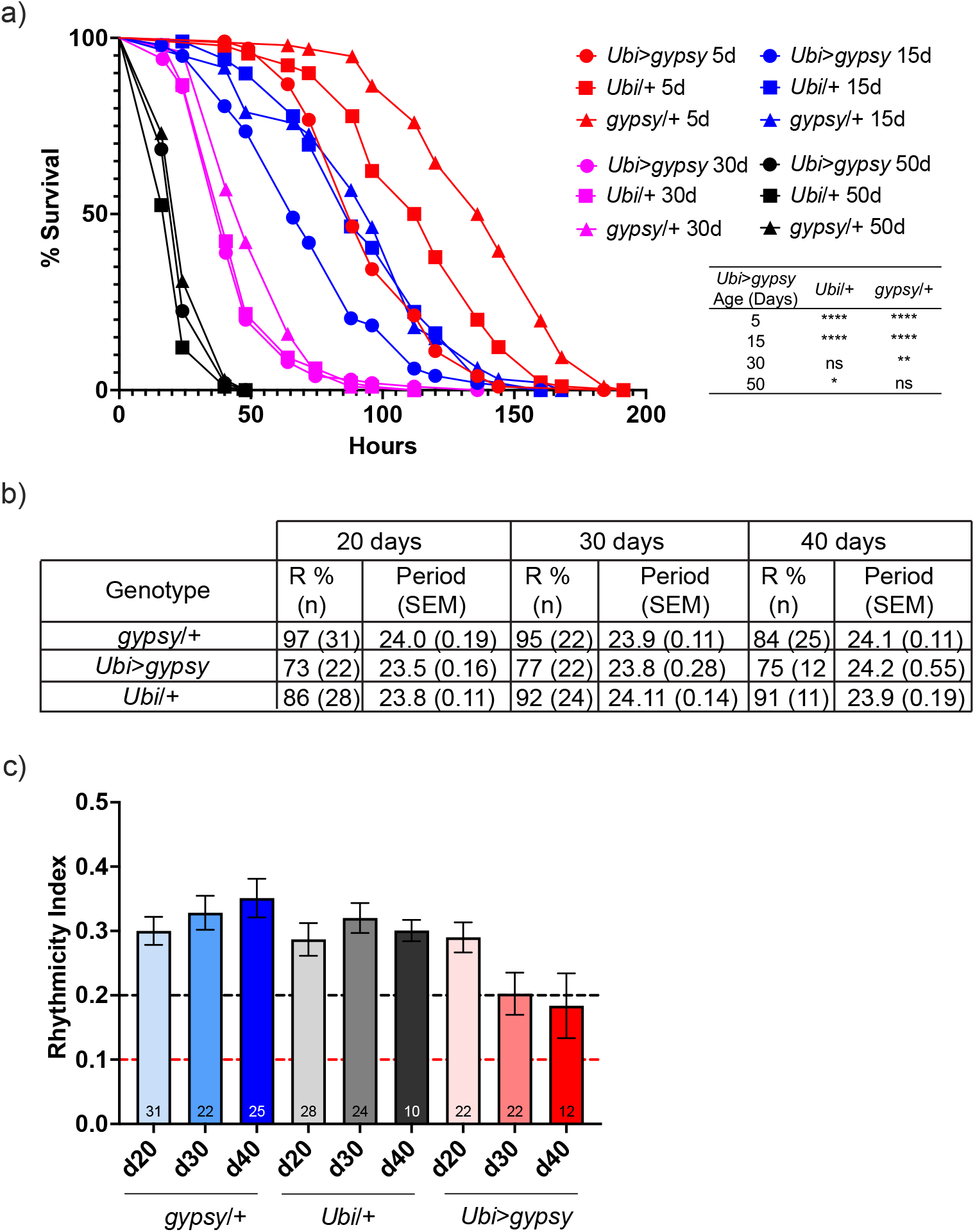
*gypsy* activity accelerates a subset of aging phenotypes. a) Survival curves of male flies after exposure to 20mM paraquat at different ages (Red = 5d, Blue = 15d, Magenta = 30d, and Black = 50d). Experimental flies expressing *gypsy* under the control of *Ubiquitin* Gal4 are represented as circles. The parental controls *Ubi*/+ and *gypsy*/+ are denoted squares and triangles, respectively. Statistical significance is indicated on graph. * p value 0.013, ** p value 0.006,**** p value <0.0001, Log-rank test. b) Summary table of the percentage of rhythmic flies (R %) and their period in DD. c) Rhythmicity levels of each genotype across the assessed ages (±SEM). *gypsy*/+ control, *Ubi*>*gypsy* experimental, and *Ubi*/+ control flies are represented in blue, red, and grey, respectively. The black and red dotted lines mark the thresholds between what are considered highly (RI>0.2) and weakly rhythmic, and arrhythmic flies (RI<0.1). The number at the bottom of each column indicates the n.

We also tested total activity as well as other well-characterized circadian phenotypes such as their ability to anticipate day and night, strength of their free-running conditions, or periodicity. Briefly, we placed 20, 30, or 40 day old flies into *Drosophila* Activity Monitors and recorded their total activity for over 10 days at 25°C in 12:12 LD conditions. We did not find any significant decrease in the overall activity levels of the active *gypsy* flies compared to their parental controls at any of the assessed times (Fig. 7-S1). Simultaneously, a separate group of flies was entrained for 3 days to the same 12:12 LD conditions and then placed in free-running conditions in constant darkness (DD) to assess the strength of the endogenous clock on these flies. While no significant effect was found in their periodicity, the rhythmicity levels of the flies overexpressing *gypsy* was affected compared to those of their age-matched controls. 84-97% of the control flies were rhythmic at all the assessed ages, whereas only 73-77% of the flies with ectopic expression of the TE remain rhythmic (Fig. 7b). Indeed, these flies also display a significant decline in their rhythmicity index throughout the assessed ages which does not happen in the control flies (Fig. 7c).

Finally, we also tested the ability of the flies with ectopic *gypsy* expression to move vertically when startled by performing a negative geotaxis assay. Briefly, we tapped the flies and recorded their location within a graduated cylinder for the following 15s. We focused on the UAS parental control to select a height and time threshold optimal across different experimental ages (Fig. 7-S2). As a result and to capture possible subtle differences between the young flies and get a robust data capture even in older ages, we focused on the percentage of flies that climbed above the 7.5 cm threshold after 5 and 10 seconds. We examined the climbing ability of the active *gypsy* flies at four different ages (7d, 14d, 35d, 56d). As expected, we observed a decay of negative geotaxis with age. However, we did not detect any accelerated decay in the *gypsy* expressing flies when compared to parental controls (Fig. 7-S3).

Having created a system in which we can regulate the activity of a particular TE (*gypsy*) and having reported the effects of its accumulation upon lifespan and other aging hallmarks, we went back to our original question regarding the role of FOXO on a specific TE. The *Ubi*-gal4 and UAS-*gypsy* lines were crossed to a UAS-*FOXO* line (Slack et al., 2011) to increase *dFOXO* expression in the animals. This leads to a 32% increase in the level of *dFOXO* mRNA in these animals (Fig. 8a). We find that increased expression of *dFOXO* can rescue the effect on lifespan due to *gypsy* activity. The 19% decrease in lifespan reported in Fig. 4a is reduced to 8% when compared to the UAS control and the lifespan matches the gal4 control (Fig. 8b). Importantly, the effect is not simply due to a prevention of ectopic *gypsy* mRNA expression as the transcript remains significantly induced in the *dFOXO* overexpressing flies (Fig. 8c).

**Figure 8.**
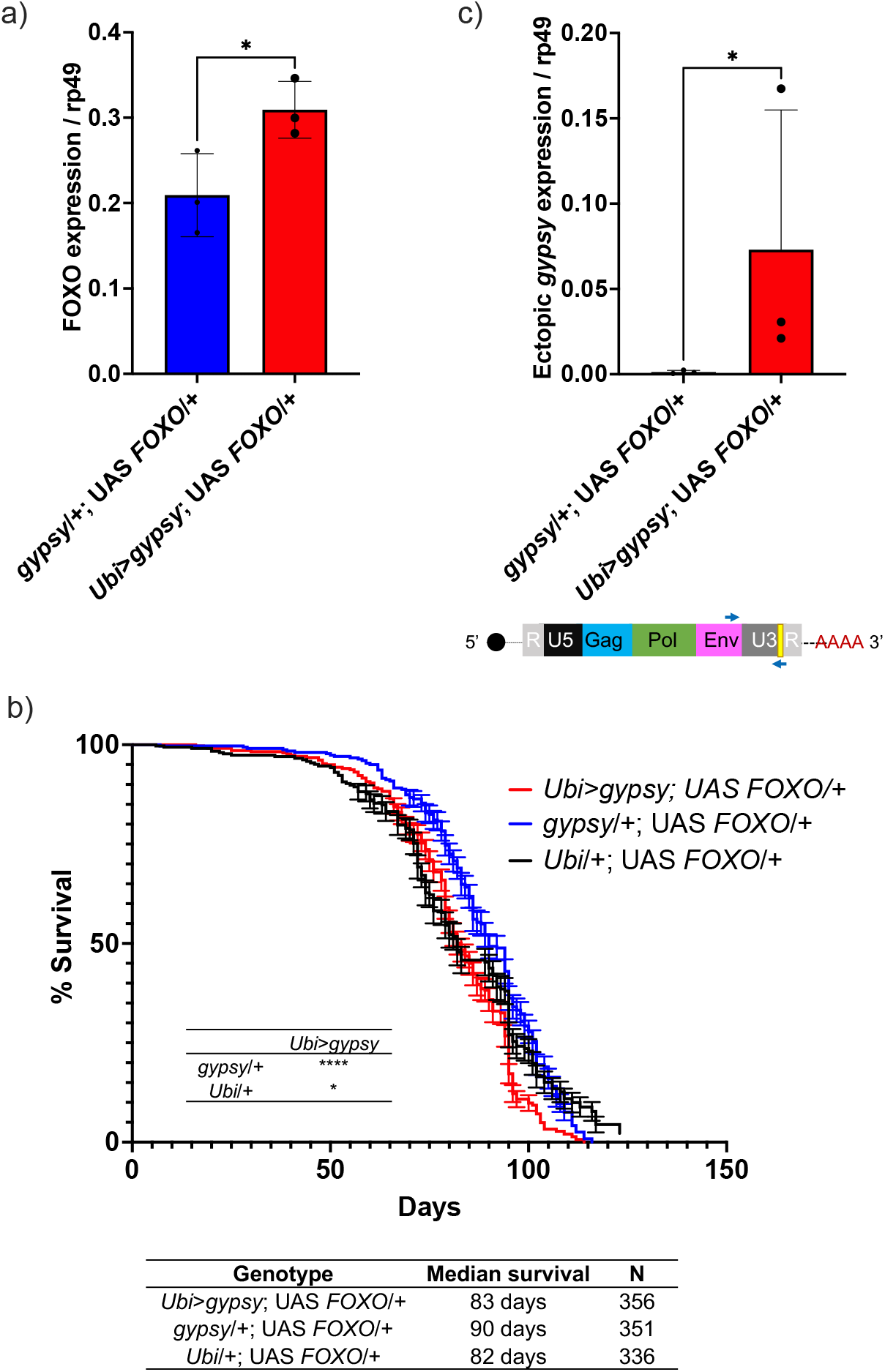
Increasing FOXO activity can rescue longevity effect. a) *dFOXO* exon 8 is detected. RT-qPCR of 5-day old males. Data are represented as means ± SD (3 biological replicates, each dot is a pool of 5 flies). One-tailed t test, * p value 0.021. b) Survival curves of male flies expressing *gypsy* and *dFOXO* under the control of *Ubiquitin* Gal4 (Red line) and the parental controls: *Ubi*/+; UAS-*FOXO*/+ (Black) and *gypsy*/+; UAS-*FOXO*/+ (Blue). Data represents 2 biological replicates, error bars SE. * 0.018 p value, **** p value <0.0001, Log-rank test. c) 3’ end of ectopic *gypsy* transcript is detected. RT-qPCR of 5-day old males. Data are represented as means ± SD (3 biological replicates, each dot is a pool of 5 flies) One tailed Mann-Whitney test, a non-parametric test, was best suited to fit the tailed distribution of the data, * p value 0.05.

## DISCUSSION

Aging can be described as a systemic breakdown due to the accumulation of different stress conditions (López-Otín et al., 2013). The stress response transcription factor FOXO can promote longevity by helping the cell respond to a myriad of conditions: oxidative stress, heat shock, virus infection and defects in protein homeostasis to name a few (Donovan & Marr, 2016; Martins et al., 2016; Spellberg & Marr, 2015). Whether FOXO can protect from the detrimental effects of TE activity on lifespan is an open question.

To begin to investigate this question we measured TE expression with age in dFOXO null and isogenic wt flies. Previous studies of wt animals report both increased and decreased TE mRNA levels with age (Chen et al., 2016; LaRocca et al., 2020). In one study of the female fatbody (5d vs 50d), researchers observed significant increases in 18 TE expression and decreases of 18 TE out of the 111 detected (Chen et al., 2016). While the total number of TE with detectable mRNA changes is lower in our whole animal study, the fact that we detect both increases and decreases in the wildtype animals is consistent with the fatbody study. In the genetic background used here, we find 18 TE have increased mRNA levels in dFOXO null flies, while only two TE showed increased mRNA in wt. Surprisingly, no TE expression is significantly decreased with age in the dFOXO null flies indicating an increase TE load in aged dFOXO null animals. Further, it suggests that the FOXO null animals are deficient in mounting a response to restrict TE expression.

Each fly strain has a unique set of TE that are capable of being transcribed. Likely due to differences in the TE landscape such as the number and location of individual TE copies in the genome (Rahman et al., 2015). We observed this difference in expressed TEs even when comparing expression in young flies (Fig. 1e.) This agrees with previous work showing supposedly isogenic stocks of *D. melanogaster* can have very different TE landscapes (Rahman et al., 2015). This difference in TE content and expression between our wt and dFOXO null strain makes it difficult to determine the effect of FOXO on individual endogenous TEs.

We created the UAS-*gypsy* system to circumvent the difference in TE landscapes and simultaneously perform a direct assay to determine whether TE activity in somatic tissue can be a causative agent of mortality and aging associated phenotypes. We chose the *gypsy* TE as our model for several reasons. First, because previous work has shown that *gypsy* insertions increase during aging, so this retrotransposon is relevant in the natural condition (Y.-H. Chang et al., 2019; Li et al., 2013). Second, a full-length clone of *gypsy* is available and it has been shown to be active and capable of transposition (Bayev et al., 1984). And lastly, the presence of only one full length copy of *gypsy* in the *D. melanogaster* reference genome (Kaminker et al., 2002) suggests a low copy number and mitigates possible unintended trans effects between the endogenous and ectopic TE. To further separate our ectopic TE, a unique 3’ sequence tag was inserted. This allows the detection and differentiation of *gypsy* mRNA and DNA content derived from our ectopic *gypsy* element.

The presence of the sequence tag in the newly formed 5’ LTR of new ectopic *gypsy* insertions allowed us to use our targeted sequencing approach to map a large number of individual new *gypsy* insertions derived from the UAS-*gypsy* synthetic TE. The target site duplication matched the site of endogenous *gypsy* elements (Dej et al., 1998) indicating that the UAS-*gypsy* element likely goes through a replication cycle identical to the natural *gypsy* element. The insertions seem to be evenly distributed and map largely to intronic sequences. This may reflect the fact that we can only recover insertions that do not have a dramatic effect on cell growth. Genomes that suffer insertions disrupting genes that are required for cell viability will be lost from the population and will not be recovered. This approach shows that the UAS-*gypsy* element can make an active transposon that can insert at sites across the genome with little chromosomal bias.

The induction of TE activity in somatic tissue resulted in a reduction in *D. melanogaster* lifespan when compared to parental controls and a significant increase in mortality in middle aged animals. There was a 19% decrease in the lifespan of male animals. Interestingly, the mortality effects of the active TE are only evident in relatively aged animals, despite the fact that the retrotransposon is active during early life. This suggests that the young animals can tolerate expression and insertion of the TE. It is only when the animals begin to age that TE expression becomes a burden and takes a toll. Perhaps it is the combination of the other metabolic and physiological effects of aging with the TE activity that is detrimental. Coincidently, this is also the timeframe that endogenous TE become expressed during normal aging (Li et al., 2013; Yang et al., 2022).

The detectable ectopic *gypsy* insertions were quantified to determine if an increase in detected insertions in the active *gypsy* flies would correlate with the decrease in lifespan. Unexpectedly, detectable insertions do not seem to increase with age, despite constant (although much more variable) expression of the transgene as the animals age. In fact, the oldest animals have the lowest detectable insertions. This finding was measured through three different approaches (the 5’ new insertion junction, the 3’ fragment of ectopic *gypsy*, and the wildtype *gypsy* env gene). All three approaches agree and show a consistent decrease with age of ectopic *gypsy* DNA content. This suggests that unknown mechanisms are acting to clear or at least prevent an increase in TE insertion load. Whether it is at the level of cellular loss or DNA repair remains to be determined.

By using a UAS-*gypsy* strain with a defective RT we were able to determine that a functional RT is needed for the TE effect on lifespan. Previous studies have suggested the need for RT to see detrimental TE effects (Gorbunova et al., 2021). The reverse transcriptase inhibitor 3TC extends the life span of a *Dcr-2* null fly strain, which has an increase in TE expression (Wood et al., 2016). The need for a functional RT to decrease lifespan implicates the DNA synthesis step as being detrimental. Because this also prevents downstream steps such as gene disruption through integration and DNA damage from incomplete integration it is not clear what step beyond DNA synthesis is most detrimental. Future experiments using this approach with integrase mutants of *gypsy* may help to answer this question.

The decrease in lifespan also opened the question of whether the active *gypsy* flies were aging more rapidly, and whether accelerated aging phenotypes could be detected. Criteria to determine premature aging and distinguish it from other causes can be summarized as follows (Salk, 2013). It must first be determined that the increase of mortality at younger ages does not alter the shape of the survival curve. An altered shape of the survival curve indicates that the health of the subjects was compromised and the increase in mortality could be due to unforeseen disease or other factors apart from natural aging processes (Piper & Partridge, 2016). Secondly, a proportional progression of all aging phenotypes without induction of disease must also be observed. The shape of the survival curve for the active *gypsy* flies indicated that the flies were healthy and thus the increase in mortality was not due to disease or external factors. The health of the active *gypsy* flies in the face of their increased mortality led us to test if aging associated phenotypes might be proportionally progressing in them. To determine if this was the case, four aging phenotypes were measured to try to detect whether an acceleration in phenotype development with respect to the parental controls was occurring in the active *gypsy* flies.

Not all phenotypes responded the same. Two out of the four phenotypes assayed show an accelerated decay. An active *gypsy* accelerates the decrease in resistance to paraquat of aging flies. Interestingly, this finding parallels the hypersensitivity to oxidative stress observed when *Dcr-2* is mutated (Lim et al., 2011). Perhaps most dramatically, 5-day old animals with an active *gypsy* have the oxidative stress resistance of a 15-day old animal. The animals’ rhythmicity is also impacted. In active *gypsy* flies, rhythmicity decays at a slightly faster rate with most animals becoming only weakly rhythmic by day 30. Other measures of circadian behavior such as total activity and period did not change with an active TE. The decay of locomotor activity also occurred at a rate similar to controls indicating that not all aging phenotypes show an accelerated decay in response to an active TE. The absence of a locomotor defect also parallels what was observed in the *Dcr-2* null fly strain (Lim et al., 2011). The distinct effects an active TE can generate on the aging phenotypes examined implies that the hallmarks of aging are not uniformly affected, and different aging processes might originate or impact a unique subset of hallmarks. We find that an active TE can accelerate a subset of aging phenotypes, and provide evidence that TE are not merely bystanders in the aging process and can behave as causative agents once they are active.

The development and use of a controllable TE expression system with a direct detrimental effect on longevity allowed us to assay whether increasing the activity of the transcription factor dFOXO played a role in promoting longevity in the face of an active TE. We find that mild overexpression of *dFOXO* can rescue the lifespan defect in the active *gypsy* flies. Though the UAS-*gypsy* was active, the decrease in lifespan was almost completely rescued, highlighting dFOXO’s ability to prolong lifespan in the face of TE activity. We and others have previously shown that FOXO responds to paraquat induced oxidative stress (Z. Chang et al., 2019; Donovan & Marr, 2016; Wang et al., 2005) and that dFOXO activates the RNAi pathway (Spellberg & Marr, 2015). Both of these responses would enhance the ability of dFOXO to combat the detrimental effect on lifespan caused by an active TE and suggests a potential new role for dFOXO in its vast repertoire to promote longevity (Martins et al., 2016).

## MATERIALS AND METHODS

### Fly stocks, *D. melanogaster* husbandry and constructs

*D. melanogaster* stocks and experimental flies were maintained at 25 °C with a 12 h light/dark cycle at controlled relative humidity. Male flies were used for this study. The fly strains used for RNA-seq were dFOXO null (*w^DAH^ ^Δ94^*) and its isogenic wildtype control *w^DAH^*, both have been previously described (Spellberg & Marr, 2015). The UAS-*gypsy* TAG fly strain was created by modifying the plasmid pDM111, which contains an active copy of *gypsy* (Bayev et al., 1984) (a generous gift from the Corces lab). The white marker from pTARG (Egli et al., 2006), and a phiC31 attB site were added to the plasmid and a unique sequence was inserted in the *gypsy* 3’LTR. The 5’ LTR of *gypsy* was precisely replaced with the UAS promoter from pUAST (Brand & Perrimon, 1993) such that the start of transcription matched the start for the *gypsy* LTR. For the RT deletion, the UAS-*gypsy* parent plasmid was cut with AflII to create an in frame deletion of most of the RT from the *gypsy* polyprotein (Marlor et al., 1986). The constructs were sent to BestGene Inc (Chino Hills, CA) for injection into *D. melanogaster*. Transgenes were integrated into the VK37 (Venken et al., 2006) attP site (BDSC 9752) using PhiC31 integrase and balanced. The line was then extensively backcrossed into the *w^1118^*background. The *Ubi*-Gal4 and UAS-FOXO fly strains were obtained from Bloomington (32551 and 42221, respectively) and crossed for at least 5 generations into the *w^1118^*lab stock. The following strains were generated by crosses: *w^1118^*; *Ubi>gypsy* (*w^1118^*; *UAS-gypsy* mated to *w^1118^*; *Ubi-Gal4*), *w^1118^*; *Ubi>ΔRT* (*w^1118^*; *ΔRT* mated to *w^1118^*; *Ubi-Gal4*), *w^1118^*; *Ubi/+*; *UAS-FOXO*/*+* (*w^1118^*; *Ubi-Gal4*;*+*/*TM3* mated to *w^1118^*; *UAS-FOXO*), *w^1118^*; *Ubi*/*UAS-gypsy*; *UAS-FOXO*/*+* and *w^1118^*; *UAS-gypsy*/+; *UAS-FOXO*/*+* (*w^1118^*; *Ubi*/*+*; *UAS-FOXO*/*+* mated to *w^1118^*; *UAS-gypsy*). All flies used throughout our experimental procedures were placed in the *w1118* genetic background.

### RNA-seq

Total RNA from the whole body of 10 male *w^DAH^*and dFOXO null (*w^DAH^ ^Δ94^*) flies was extracted with TRI Reagent at 5 to 6 days and 30 to 31 days old according to the manufacturers protocol (Molecular Research Center, Inc., Cincinnati, OH). To generate RNA-seq libraries, 1 µg of total RNA was used as input for the TruSeq RNA Library Prep Kit v2 (Illumina, Inc., San Diego, CA) and the manufacturer’s protocol was followed. Libraries were sequenced on an Illumina NextSeq 500 in 1 x 75 bp mode. Three individually isolated biological replicates were sequenced for each condition.

### Bioinformatics

RNA-seq fastq files were uploaded to the public server usegalaxy.org and processed at the Galaxy web platform (Afgan et al., 2018). The tools FASTQ Groomer (Blankenberg et al., 2010) and FastQC (Andrews, 2010) were used for library quality control. To obtain gene counts, the RNA STAR aligner (Dobin et al., 2012) was used to map the sequencing data to both the *D. melanogaster* genome (dm6) and to a TE consensus FASTA file with 176 *Drosophila* TE. The R package DEBrowser (Kucukural et al., 2019) was used for the following procedures: to filter the counts (1 count per million (CPM) in at least 11/12 libraries), 105 TE passed filtering, calculate differential expression and statistical significance with DESeq2 (parameters: 5% false discovery, 1.5 fold, local, no beta prior, LRT), and generate volcano and scatter plots.

### Life Span Assays

Life span assays were performed as previously described (Linford et al., 2013). Briefly, newly eclosed flies were mated for 48hr, sorted by gender, and kept at a standard density of 15 flies per vial. Flies were transferred to fresh food every 2-3 days and mortality was recorded. 3 independent biological replicates of at least 100 flies were performed at different times for *Ubi*>*gypsy*. The age specific mortality rate was calculated by dividing the deaths occurred in a given age group by the size of the population in which the deaths occurred (*Principles of Epidemiology | Lesson 3 - Section 3*, 2021). 2 independent biological replicates were performed at different times for *Ubi*>*gypsy*; UAS-FOXO/+ and *Ubi>ΔRT*. Kaplan-Meier survival curves were generated with GraphPad Prism version 9 (GraphPad Software, San Diego, California USA, www.graphpad.com) and analyzed with the Log Rank test.

### RT-qPCR and genomic DNA qPCR

Total RNA or DNA was extracted from the whole body of 5-10 male flies. RNA was extracted with the TRI Reagent according to the manufacturers protocol (Molecular Research Center, Inc., Cincinnati, OH) and genomic DNA as described previously (Aljanabi & Martinez, 1997). cDNA for RT-qPCR was synthesized as previously described (Olson et al., 2013). RT-qPCR and gDNA qPCR were performed as follows: for a 10 µL reaction 2 µL of cDNA or 50ng of DNA were used as template and assayed with SYBR green with the primers in Table 1. For all experiments, three biological replicates were assayed for each condition and the relative expression was calculated as a fraction of the housekeeping gene Rp49.

**Table 1.**
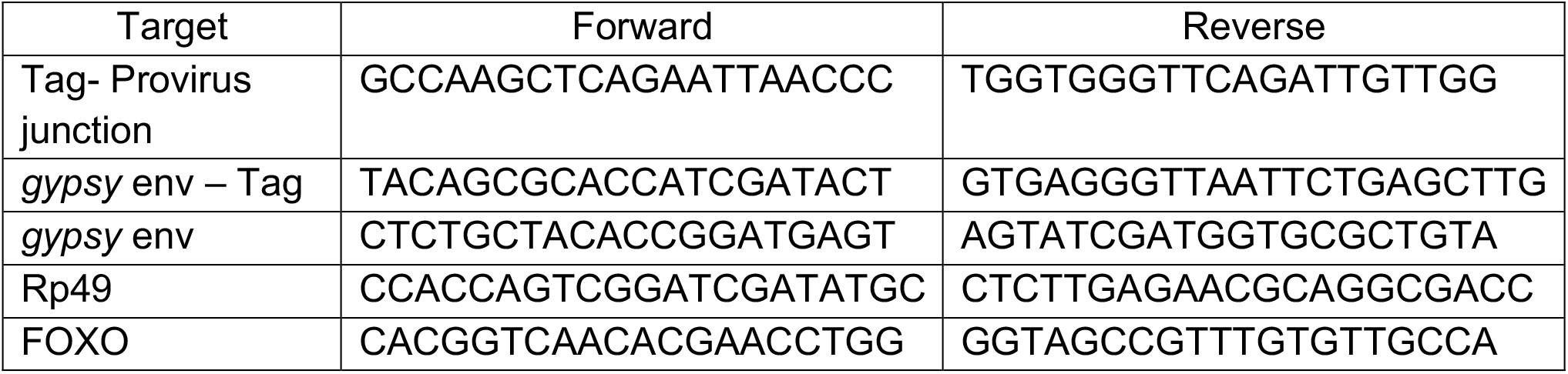
qPCR Primers.

**Table 2.**
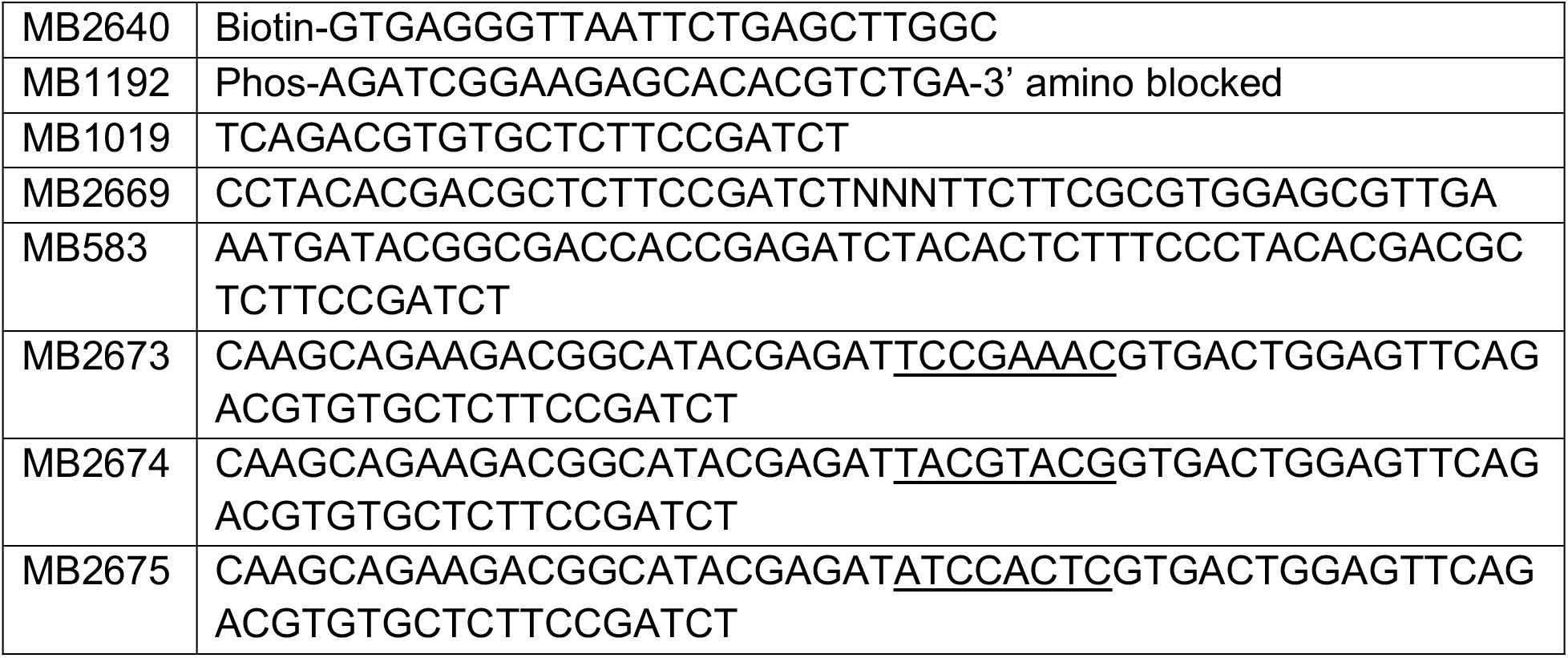
Primers for NGS sequencing.

### Oligo adenylation

MB1192 was adenylated using recombinant MTH ligase (Zhelkovsky & McReynolds, 2011). 200pmol of oligo was adenylated in a 200µl reaction (50mM Tris 7.5, 10mM MgCl2, 0.1mM EDTA, 0.1mM ATP, 5mM DTT) with 10µl recombinant MTH ligase. The reaction was incubated at 65°C for 1 hour and terminated by incubation at 85°C for 15 minutes. 20µg of glycogen was added to the reaction followed by 500µl ethanol. The oligo was placed at −20°C for 30 minutes and collected by centrifugation. The pellet was resuspended in 100µl 6M GuHCl, 50mM Tris 6.8. 400µl of ethanol was added and the oligo was loaded onto a silica spin column (BioBasic PCR cleanup). The column was washed once with 80% ethanol and then centrifuged until dry. The adenylated oligo was eluted in 40µl TE and quantitated by UV absorbance.

### NGS library preparation

To prepare the insertion libraries, 250ng of genomic DNA was digested with the 4-base cutter MnlI (NEB, Ipswich, MA), overnight at 37°C according to manufacturer’s recommendations, to fragment the genome. Additionally, MnlI cuts the UAS-*gypsy* element 61 times including 17bp from the 5’ end of the 3’LTR to limit the recovery of the parental UAS-*gypsy* sequences. The fragmented DNA was purified using a silica-based PCR cleanup kit (Biobasic, Markham, Canada). A biotinylated primer (MB2640) annealing to the TAG sequence was annealed and extended with 20 cycles of linear amplification with pfuX7(Nørholm, 2010) in a 100µl reaction (20mM Tris-HCl pH 8.8 at 25°C, 10mM (NH4)2SO4, 10mM KCl, 2mM MgSO4, 0.1% Triton® X-100, 200µM dNTPs, 0.5µM primer, 5u pfuX7) The reaction was quenched by adding 400µl TENI (10mM Tris 8.0, 1mM EDTA, 25mM NaCl, 0.01% Igepal 630). Biotinylated products were purified using M270 dynabeads (ThermoFisher, Waltham, MA) by incubating the reaction for 30 minutes at room temperature followed by magnetic separation of bound DNA. The beads were washed three times with TENI. Single-stranded biotinylated DNA was purified by incubating the beads with 0.15N NaOH for 15 minutes at room temperature. Beads were washed once more with 0.15N NaOH to remove the non-biotinylated DNA strand. Beads were neutralized by washing two times in TENI and transferred to a new tube. The adenylated MB1192 oligo was ligated to the biotinylated ssDNA on the dynabeads using MTH ligase(Torchia et al., 2008). Beads were resuspended in a 60µl reaction containing 10mM Hepes pH7.4, 5mM MnCl2 and 60pmol adenylated MB1192 oligo. 5µl recombinant MTH RNA ligase was added and the reaction was incubated at 65°C for 1 hour and terminated by incubation at 85°C for 15 minutes. The library was amplified by PCR using a primer to the ligated product (MB1019) and a nested primer containing a 5’ tail with illumina sequences (MB2669). The individual libraries were amplified by PCR using primers containing attachment sequences (MB583) and barcodes (MB26673 or MB2674 or MB2675). Libraries were sequenced on an illumina Miseq using a PE 150 kit.

### Insertion mapping

To process the reads, the fastQ file was first separated by illumina barcode. The individual libraries were processed using Galaxy (Afgan et al., 2018). First MB2667 primer sequences were removed from read 1 using Trimmomatic (Bolger et al., 2014). Reads containing the *gypsy* LTR were separated using Barcode Splitter. The remaining LTR sequences were removed using cutadapt (Martin, 2011). The reads were filtered for size and then aligned to the UAS-*gypsy* construct. 20-35% of the reads mapped to the original parental UAS-*gypsy*. Unaligned reads were then aligned to the *Drosophila* genome (dm6) using Bowtie2 (Langmead & Salzberg, 2012). PCR duplicates were removed, and the insertion sites were compared to the flybase genespan annotation to identify all insertions in transcribed regions. Insertion sites were compared to the intron annotation for Ensembl for identifying insertions in introns. The reverse complement of the first 22 nucleotides of the deduplicated insertion sites were used to determine the probability of finding each nucleotide at each position using weblogo3 (Crooks et al., 2004).

### Paraquat stress assay

Male flies generated and reared in the same conditions as life span assay flies were fed 20mM paraquat at 5, 15, 30, or 50 days. Briefly, at the specified time point flies were starved for 3-4 hr prior to transfer into a minimal food medium (5% sucrose, 2% agar, water) containing 20mM paraquat. Mortality was recorded in 8 and 16 hr intervals. 95-100 flies were used per genotype/timepoint for all time points except 5 day Ubi/+ which had 90 flies. Kaplan-Meier survival curves were generated with GraphPad Prism version 9 (GraphPad Software, San Diego, California USA) and analyzed with the Log Rank test.

### Negative Geotaxis Assay

A negative geotaxis assay based on previously established protocols (Gargano et al., 2005; Madabattula et al., 2015; Tuxworth et al., 2019) was set up to assay experimental and control flies at 7, 14, 35, and 56 days. Briefly, male flies generated and reared in life span assay conditions were used. The same cohort was assayed for all the different time points. The day before the experiment, flies were transferred under light CO2 to fresh food vials (10 flies per vial) and allowed to recover for at least 18 hr. Flies were then taken out of the incubator and allowed to acclimate to room temperature for 1 hr. They were then transferred to a 50 mL graduated glass cylinder (VWR, Radnor, PA) sealed with a cotton plug and allowed to acclimate for 1 min before starting trials. Trials were recorded on an iPhone X camera (Apple Inc, Cupertino, California) placed 30 cm away from the recording spot. A trial consisted in tapping flies to the base of the cylinder and allowing them to climb for 20 seconds. They were then given 60 seconds to recover before starting the next trial. A total of 5 trials was performed for each vial. 10 vials were assayed per genotype. N = 100. Flies were then transferred to fresh food vials and returned to the incubator. Linear regression was performed in Prism (GraphPad Software, San Diego, California USA, www.graphpad.com) to determine the slope of the curves and any possible significant differences.

### Locomotor Activity Analysis

Flies were entrained in 12:12 LD (light-dark) conditions at 25 °C. The locomotor activity of 20, 30, and 40 days old male flies was recorded using the Trikinetics locomotor activity monitor (Waltham, MA). Two sets of experiments were conducted. On one set, flies were maintained in 12:12 LD throughout the whole length of the experiment (10-12 days). The other set, was entrained in 12:12 for 3 days, followed by at least 5 days in constant darkness (DD). Quantification of total activity and the analysis of circadian rhythmicity strength and period were conducted in Matlab using Vecsey’s SCAMP (Donelson et al., 2012). Flies with a rhythmicity index (RI) < 0.1 were classified as arrhythmic; ones with RI 0.1-0.2, as weakly rhythmic, while flies with RI >0.2 were considered strongly rhythmic. Only the period of weakly or strongly rhythmic flies were included in the calculation of the free-running period for each genotype. N is included in figure.

### Statistics

All statistical analyses, except RNA-seq, were conducted using GraphPad Prism version 9 for Mac (GraphPad Software, San Diego, California USA, www.graphpad.com). Multiple comparisons in the Ordinary one-way and 2way ANOVA were corrected by the two-stage linear step-up procedure of Benjamini, Krieger and Yekutiel with a false discovery rate of 5%.

### Materials availability statement

The transgenic fly strains created during this study can be obtained by request to the corresponding author. All next generation sequencing data generated are undergoing submission at the GEO database and will be publicly accessible.

## SUPPLEMENTS

**Figure 1-S1.**
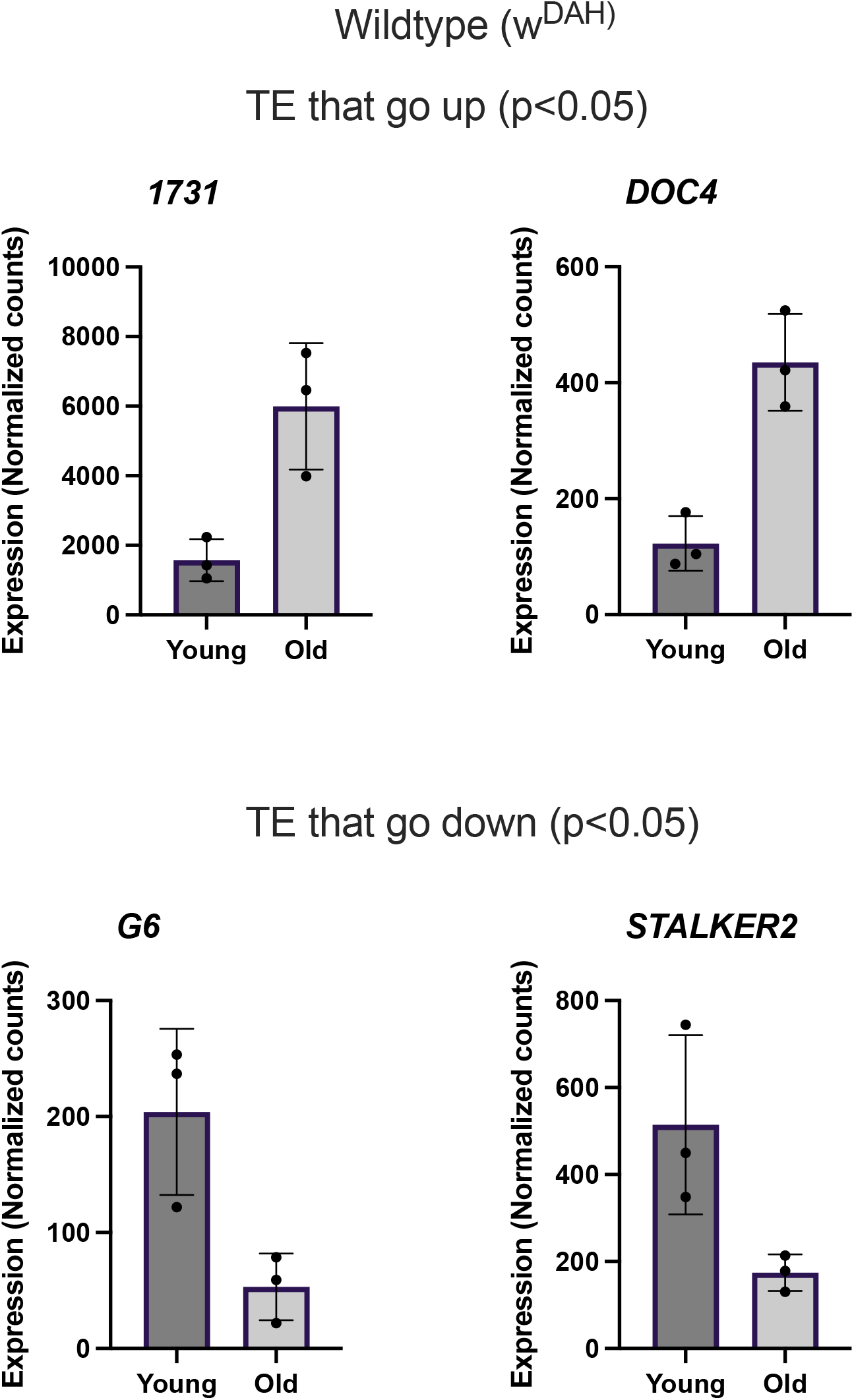
Differentially expressed TE in wild type. Median ratio normalized counts by DEseq2 of TE differentially expressed with age in wt flies.

**Figure 1-S2.**
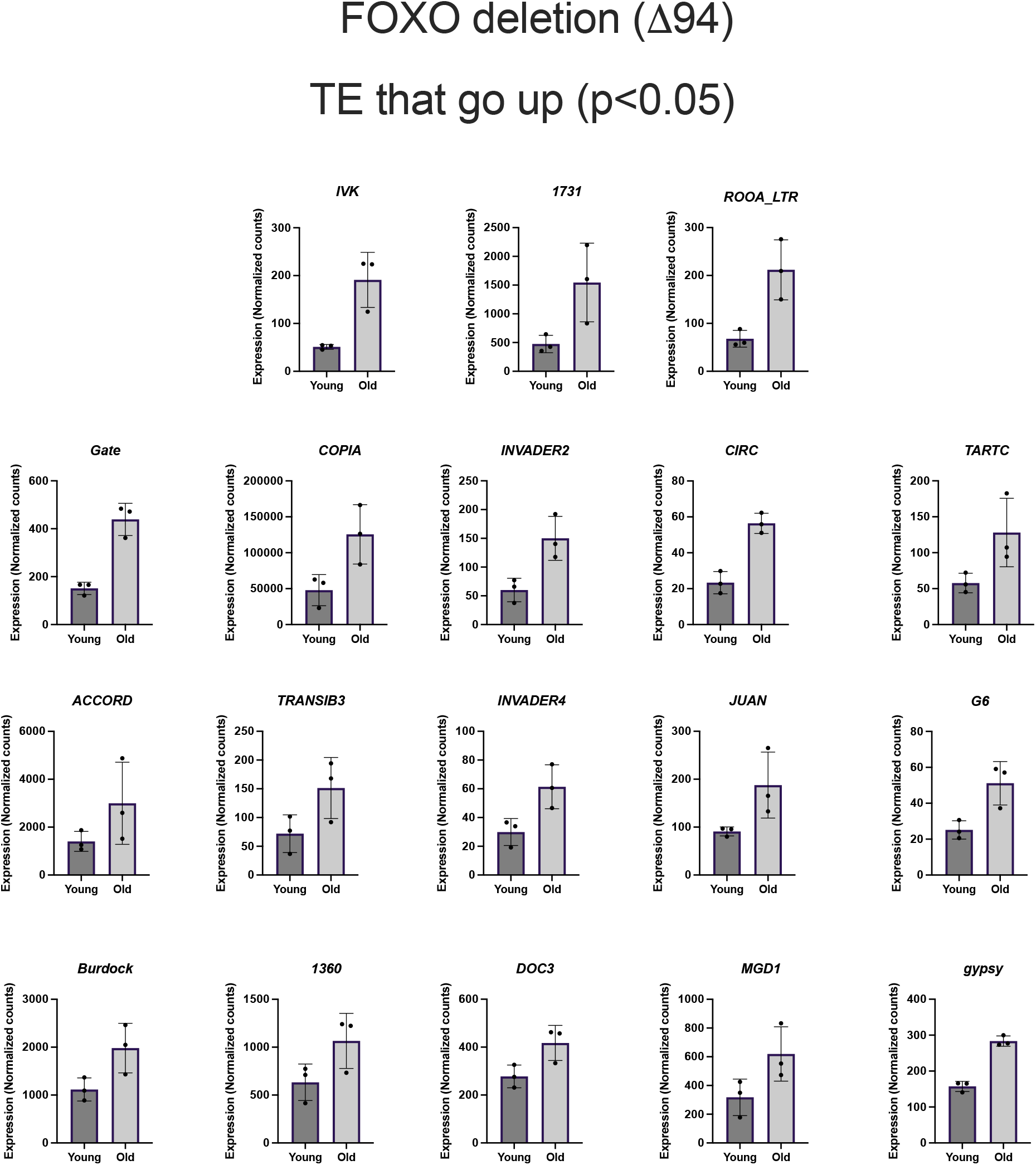
Differentially expressed TE in FOXO deletion flies. Median ratio normalized counts by DEseq2 of TE differentially expressed with age in FOXO deletion flies.

**Figure 3-S1.**
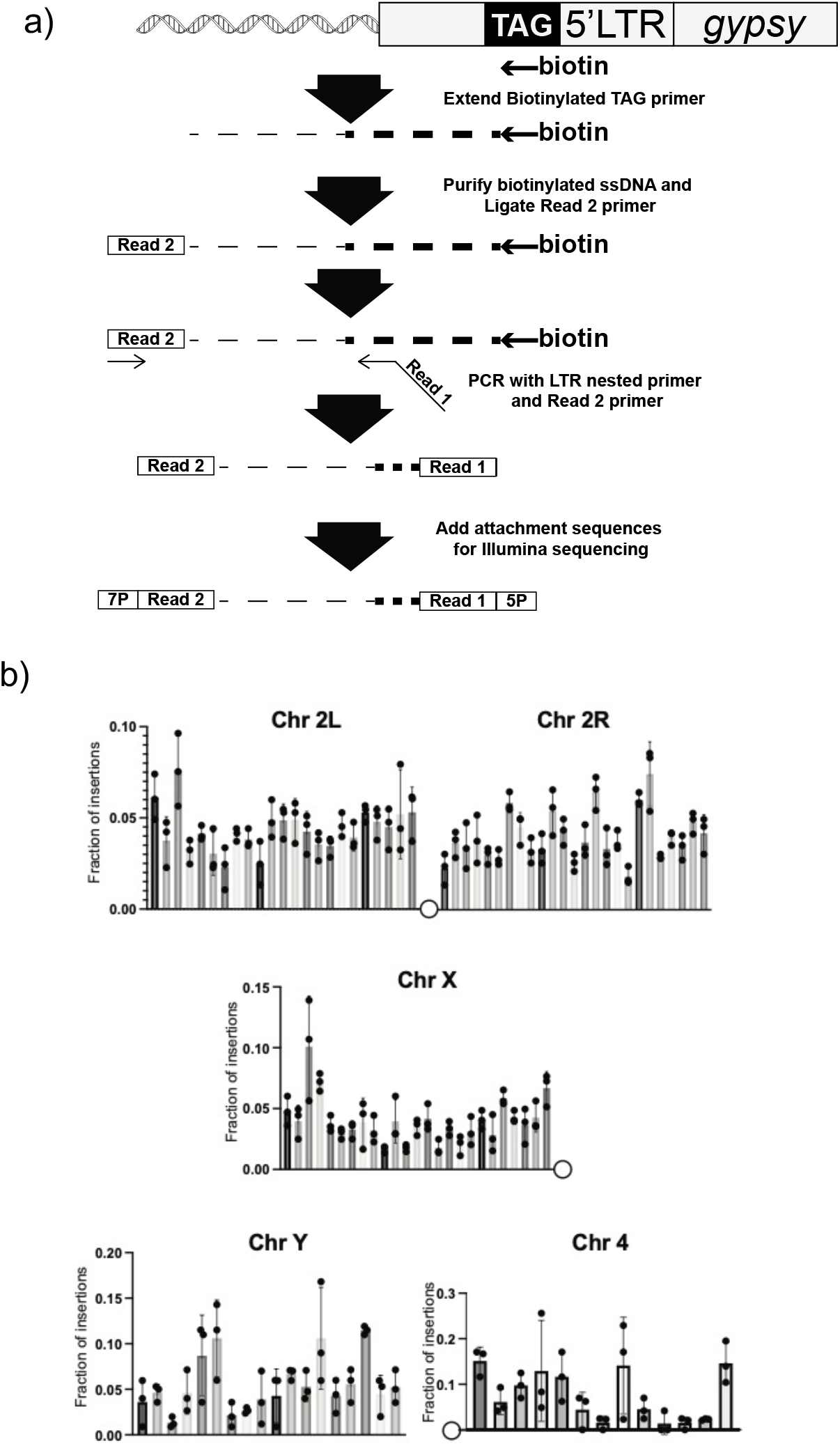
NGS mapping of ectopic *gypsy* insertions. a) Schematic of targeted sequencing approach to map 5’ junctions. b) The fraction of insertions that map to each one megabase region of the reference genome for the arms of chromosome 2 and the X chromosome are plotted. For the Y chromosome, each bin represents 200K bases. For the chromosome 4, each bin represents 100K bases. For all histograms, the bars represent the average of three biological replicates. Error bars indicate the standard deviation and the filled circles indicate the individual measurements.

**Figure 4-S1.**
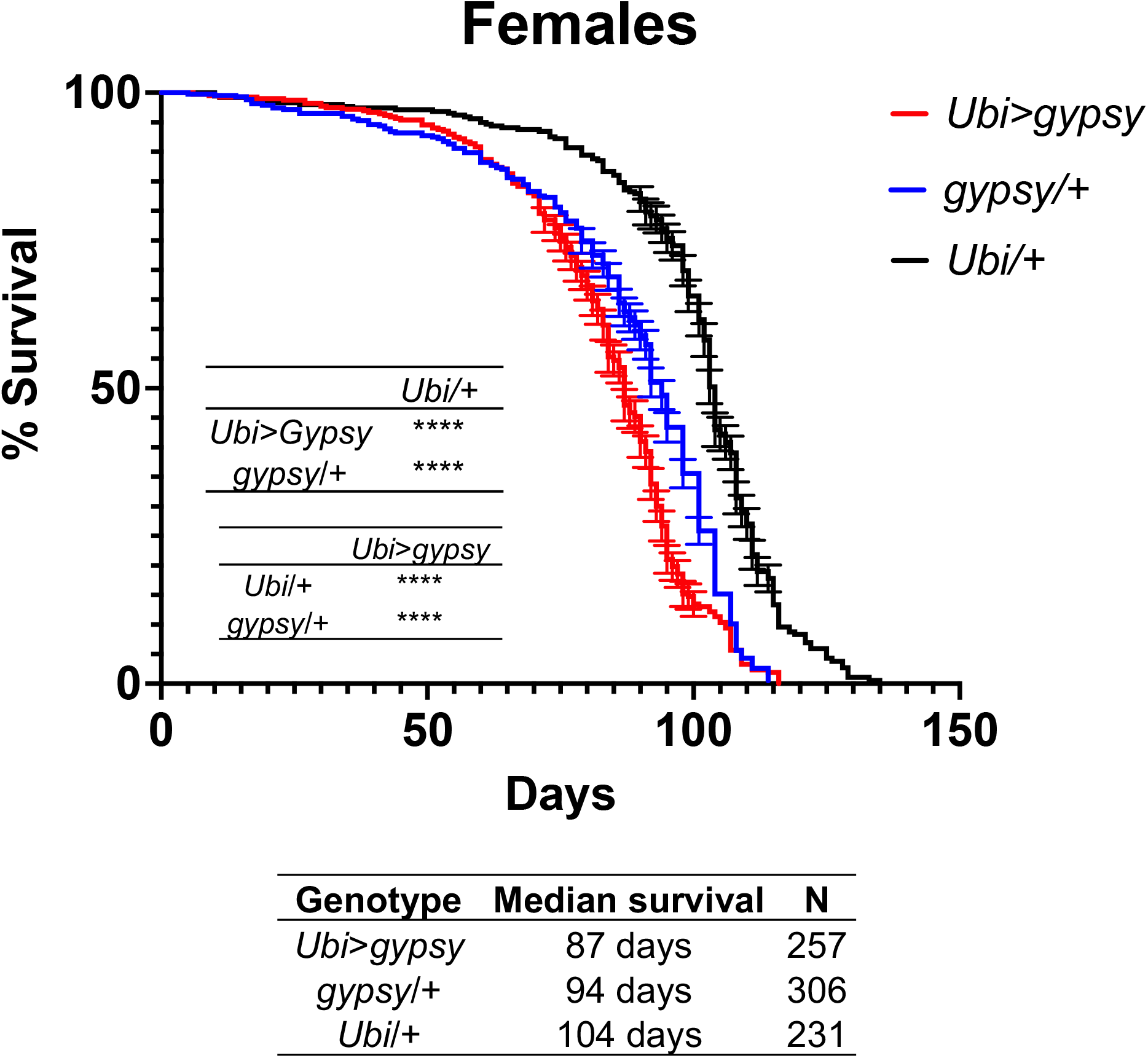
Female parental UAS-*gypsy* control has shortened lifespan. a) Survival curves of female flies expressing *gypsy* under the control of *Ubiquitin* Gal4 (Red) and the parental controls: *gypsy*/+ (Blue) and *Ubi*/+ (Black). Data represents 3 biological replicates (independent cohorts done at different times of year), error bars SE. **** p value <0.0001, Log-rank test.

**Figure 4-S2.**
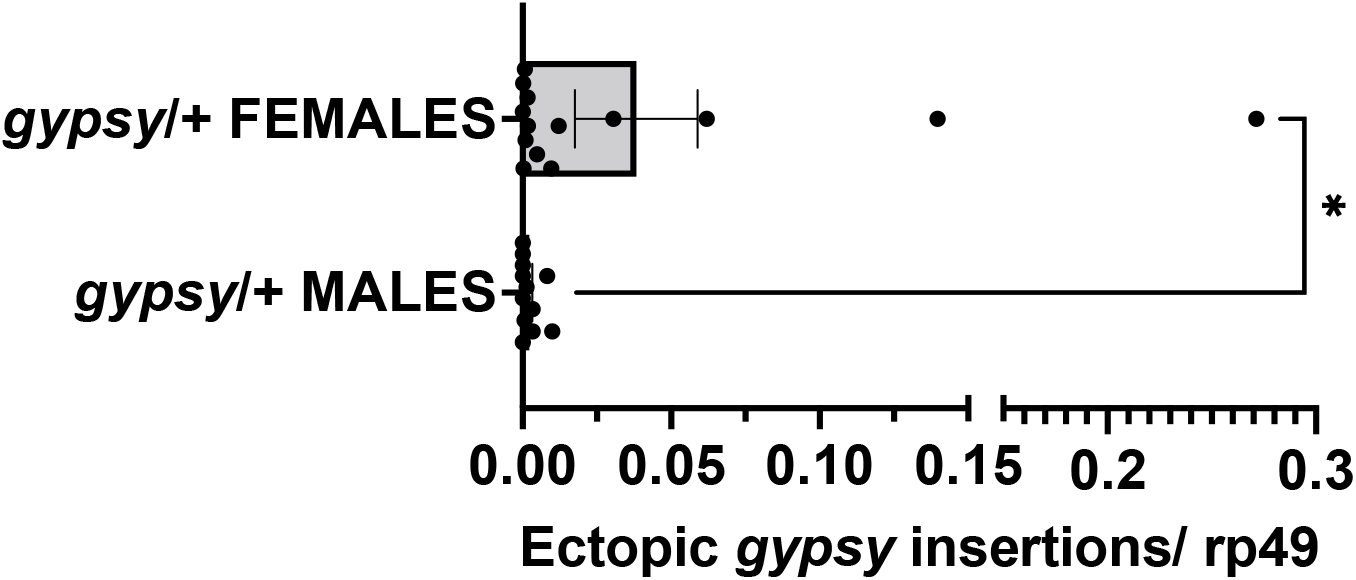
Male vs female parental UAS *gypsy* control. Ectopic *gypsy* provirus insertion junctions are detected. Data are represented as means ± SEM. *gypsy*/+ gDNA from dead flies across the whole lifespan curve was assayed. All dead flies were assayed. Each dot represents a pool of 5-10 dead flies. Two tailed Mann-Whitney test, * p value 0.0177. Male n=12 pools. Female n = 14 pools.

**Figure 4-S3.**
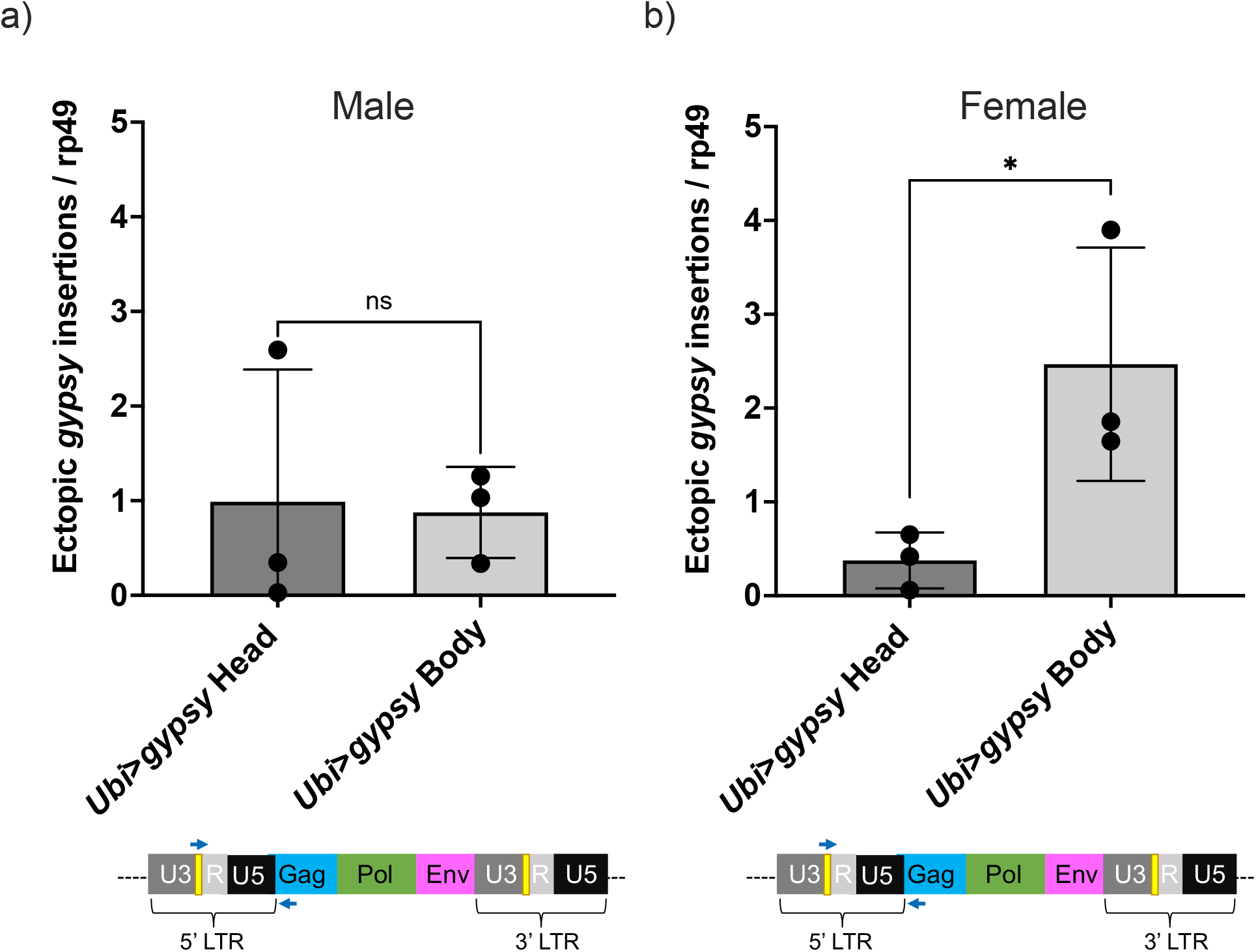
Head vs body *gypsy* parental control. a-b) Ectopic *gypsy* provirus insertion junctions are detected. Data are represented as means ± SD (3 biological replicates, each dot represents a pool of gDNA from 15-20 14d-old heads or bodies). a) Males. 2 tailed unpaired t test, ns p value >0.9. b) Females. 2 tailed unpaired t test, * p value 0.047.

**Figure 7-S1.**
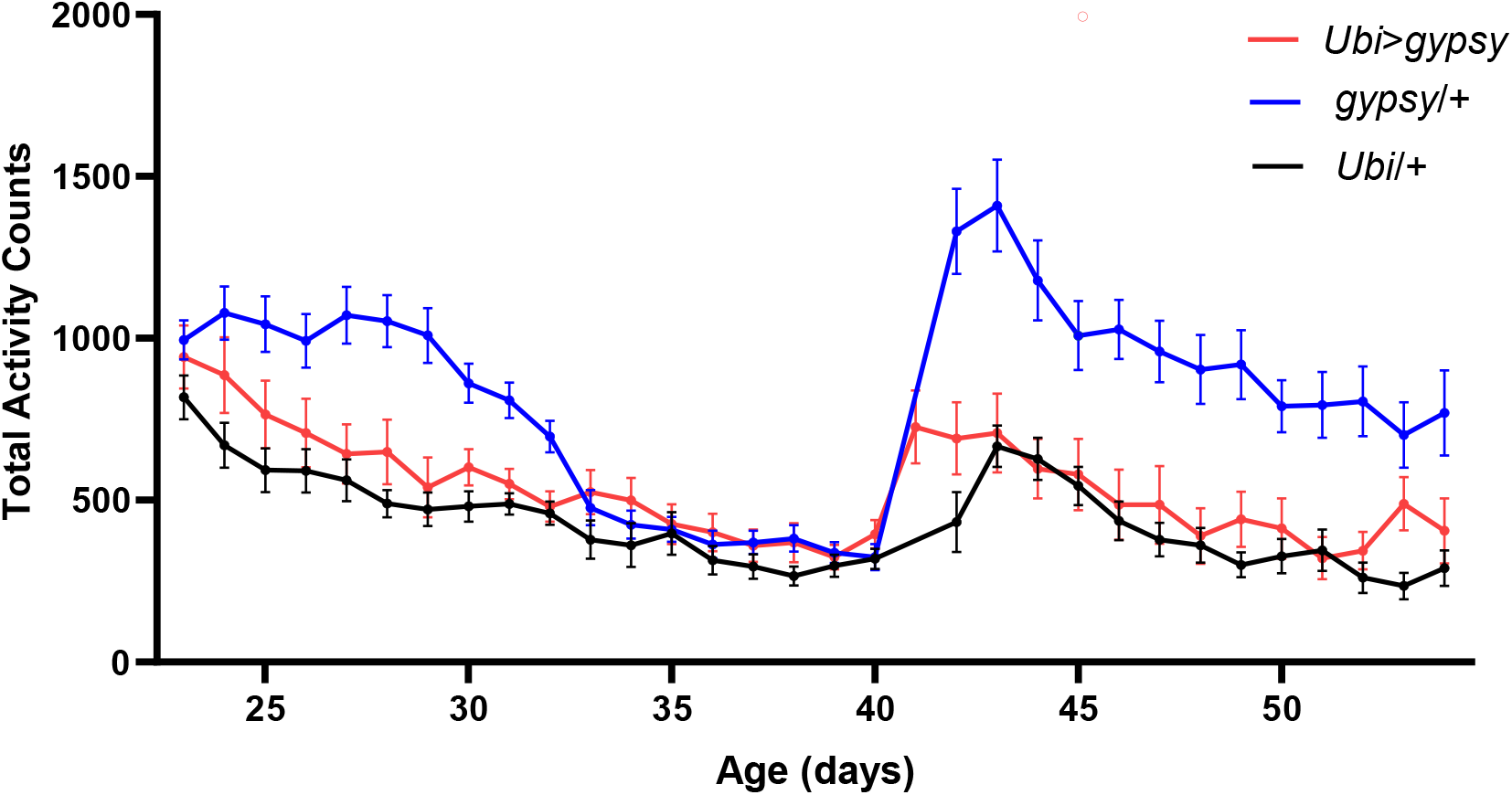
Total activity counts of each genotype *per* day under 12:12 LD conditions. The sudden increase in activity around day 40 corresponds with the start of a new recording. UAS-*gypsy*/+ blue line. *Ubi*>*gypsy* red line. *Ubi*/+ black line. N=26-32. Data are represented as means ± SEM.

**Figure 7 S2.**
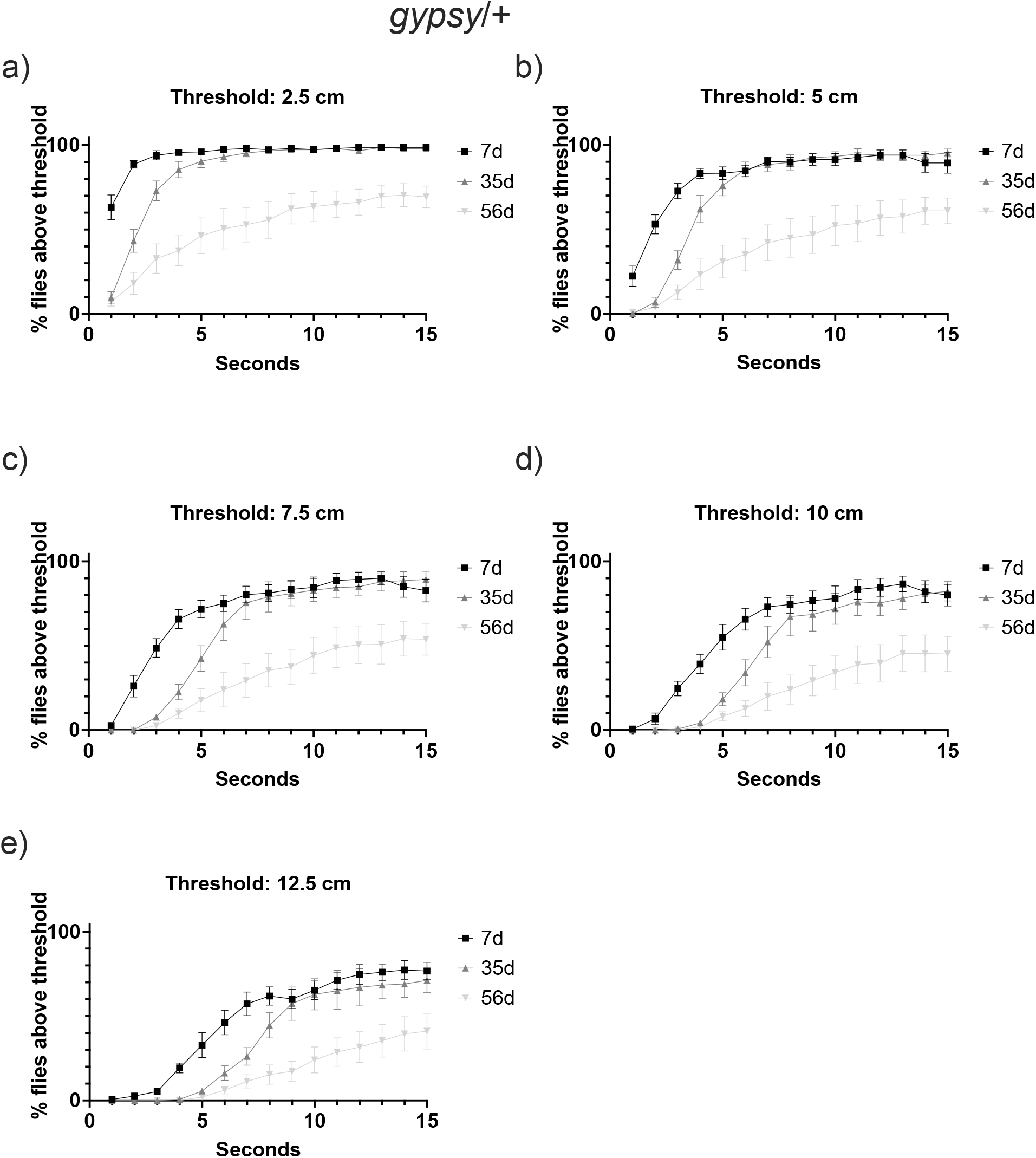
Negative geotaxis threshold optimization. Climbing behavior at 3 different ages (7 day old (black square), 35 day old (grey triangle), and 56 day old (light grey inverted triangle) of the UAS-*gypsy* parental control. Data are represented as means ± SEM (5 biological replicates, 3 trials each).

**Figure 7-S3.**
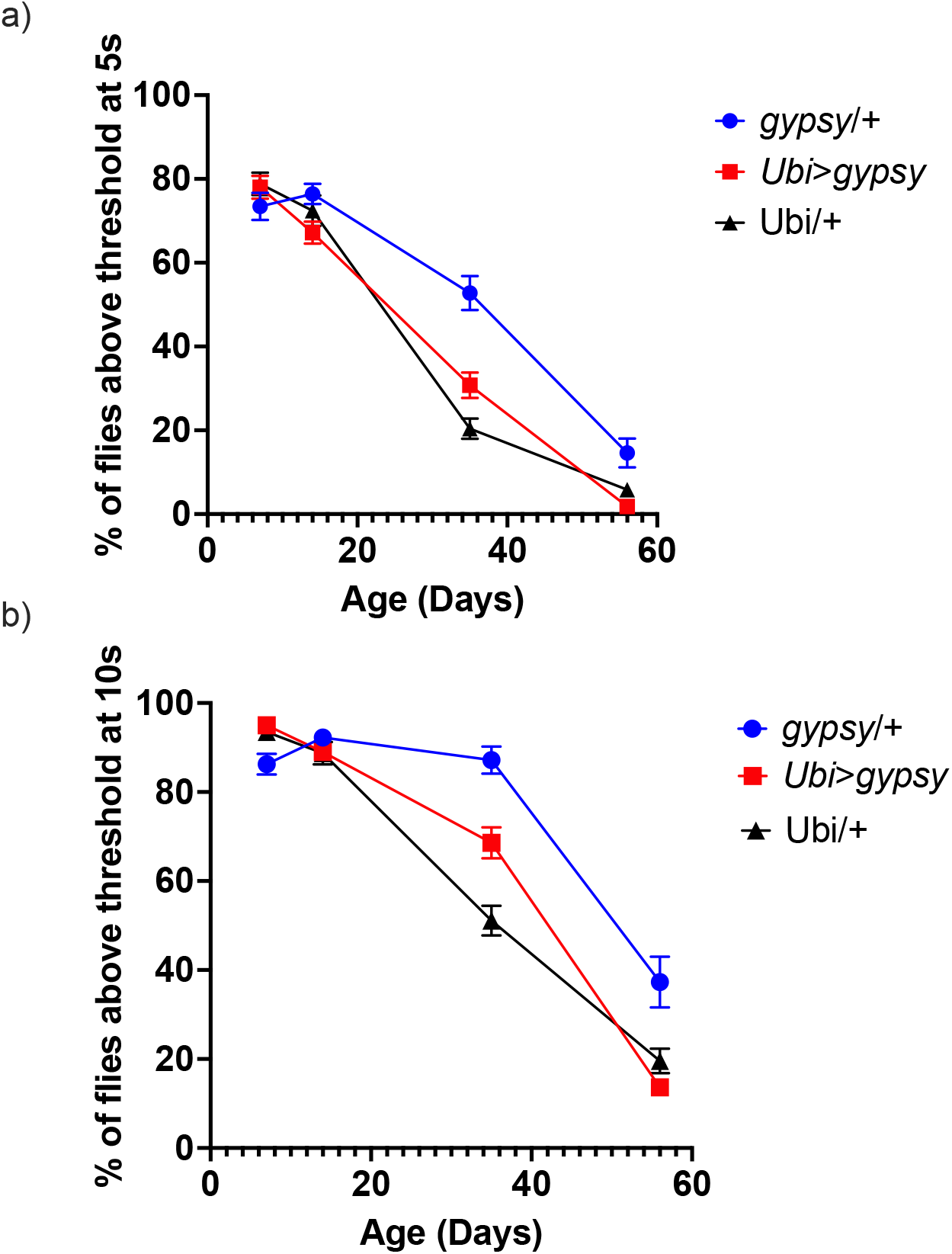
Negative geotaxis. Threshold 7.5 cm. Data are represented as means ± SEM (10 biological replicates, 10 male flies each, 5 trials per replicate). *gypsy*/+ blue circles. *Ubi*>*gypsy* red squares. *Ubi*/+ black triangles. A) 5 seconds. B) 10 seconds. Slopes were analyzed by simple linear regression and determined to be not significantly different from each other.

**Figure 1-Table supplement 1 (WT).** Wild type control TE expression. Counts have been normalized by median ratio normalization in DESeq2. Adjusted p value calculated by DEseq2 is provided.

| TE | wt_6d_1 | wt_6d_2 | wt_5d_3 | wt_30d_1 | wt_30d_2 | wt_30d_3 | adjusted p value | Fold Change | Class |
| --- | --- | --- | --- | --- | --- | --- | --- | --- | --- |
| **17.6** | 57 | 108 | 49 | 29 | 100 | 98 | 9.97E-01 | 1.06 | Retrotransposon |
| **297** | 722 | 1071 | 787 | 68 | 409 | 684 | 9.39E-01 | 0.45 | Retrotransposon |
| **412** | 136 | 174 | 95 | 61 | 126 | 238 | 9.97E-01 | 1.05 | Retrotransposon |
| **1731** | 1432 | 2236 | 1051 | 3987 | 6466 | 7530 | 2.10E-04 | 3.81 | Retrotransposon |
| **ACCORD** | 49 | 436 | 21 | 1 | 40 | 157 | 9.97E-01 | 0.39 | Retrotransposon |
| **aurora-element** | 41 | 27 | 27 | 35 | 29 | 28 | 9.97E-01 | 0.98 | Retrotransposon |
| **BAGGINS** | 28 | 43 | 19 | 12 | 31 | 60 | 9.97E-01 | 1.14 | Retrotransposon |
| **Beagle** | 486 | 391 | 455 | 343 | 400 | 369 | 9.97E-01 | 0.83 | Retrotransposon |
| **Beagle2** | 23 | 56 | 11 | 11 | 25 | 54 | 9.97E-01 | 0.99 | Retrotransposon |
| **Bel** | 1000 | 1365 | 314 | 254 | 882 | 1594 | 9.97E-01 | 1.02 | Retrotransposon |
| **BLOOD** | 488 | 578 | 505 | 135 | 499 | 381 | 9.97E-01 | 0.65 | Retrotransposon |
| **BS** | 63 | 84 | 27 | 26 | 49 | 66 | 9.97E-01 | 0.81 | Retrotransposon |
| **BS3** | 66 | 60 | 23 | 65 | 59 | 52 | 9.97E-01 | 1.17 | Retrotransposon |
| **Burdock** | 1044 | 1138 | 704 | 900 | 1811 | 1675 | 9.96E-01 | 1.52 | Retrotransposon |
| **CIRC** | 43 | 41 | 17 | 14 | 45 | 73 | 9.97E-01 | 1.30 | Retrotransposon |
| **COPIA** | 78001 | 138845 | 60568 | 95444 | 238381 | 250318 | 6.22E-01 | 2.11 | Retrotransposon |
| **CR1A** | 333 | 463 | 124 | 142 | 344 | 419 | 9.97E-01 | 0.98 | Retrotransposon |
| **Diver** | 131 | 127 | 19 | 7 | 71 | 121 | 9.97E-01 | 0.72 | Retrotransposon |
| **DIVER2** | 62 | 51 | 10 | 4 | 33 | 46 | 9.97E-01 | 0.68 | Retrotransposon |
| **DM_ROO** | 1320 | 1461 | 521 | 898 | 1373 | 1649 | 9.97E-01 | 1.19 | Retrotransposon |
| **DM88** | 57 | 62 | 16 | 22 | 50 | 66 | 9.97E-01 | 1.02 | Retrotransposon |
| **DMGYPF1A** | 777 | 984 | 454 | 311 | 644 | 862 | 9.97E-01 | 0.82 | Retrotransposon |
| **DOC** | 2501 | 4171 | 1162 | 480 | 2344 | 3052 | 9.97E-01 | 0.75 | Retrotransposon |
| **DOC2** | 241 | 303 | 40 | 52 | 151 | 248 | 9.97E-01 | 0.77 | Retrotransposon |
| **DOC3** | 114 | 134 | 63 | 84 | 146 | 138 | 9.97E-01 | 1.18 | Retrotransposon |
| **DOC4** | 105 | 177 | 88 | 525 | 422 | 359 | 1.00E-05 | 3.52 | Retrotransposon |
| **DOC5** | 8 | 18 | 9 | 14 | 26 | 36 | 1.00E+00 | 2.14 | Retrotransposon |
| **F-element** | 392 | 646 | 138 | 74 | 304 | 594 | 9.97E-01 | 0.83 | Retrotransposon |
| **Flea** | 1122 | 1506 | 397 | 310 | 1042 | 1294 | 9.97E-01 | 0.87 | Retrotransposon |
| **FROGGER** | 59 | 62 | 6 | 7 | 42 | 57 | 9.97E-01 | 0.84 | Retrotransposon |
| **FW2** | 38 | 44 | 15 | 19 | 45 | 41 | 9.97E-01 | 1.08 | Retrotransposon |
| **FW3** | 43 | 38 | 27 | 34 | 36 | 42 | 9.97E-01 | 1.04 | Retrotransposon |
| **G2** | 87 | 71 | 33 | 28 | 56 | 73 | 9.97E-01 | 0.82 | Retrotransposon |
| **G3** | 15 | 21 | 12 | 4 | 22 | 23 | 1.00E+00 | 1.02 | Retrotransposon |
| **G4** | 129 | 142 | 74 | 73 | 130 | 160 | 9.97E-01 | 1.05 | Retrotransposon |
| **G5** | 57 | 74 | 11 | 27 | 70 | 72 | 9.97E-01 | 1.19 | Retrotransposon |
| **G5A** | 184 | 185 | 155 | 147 | 175 | 218 | 9.97E-01 | 1.03 | Retrotransposon |
| **G6** | 122 | 237 | 253 | 22 | 79 | 59 | 8.30E-03 | 0.26 | Retrotransposon |
| **Gate** | 252 | 288 | 69 | 240 | 563 | 881 | 3.95E-01 | 2.76 | Retrotransposon |
| **GTWIN** | 28 | 78 | 13 | 36 | 70 | 62 | 9.97E-01 | 1.40 | Retrotransposon |
| **GYPSY10** | 179 | 225 | 102 | 110 | 197 | 228 | 9.97E-01 | 1.06 | Retrotransposon |
| **GYPSY11** | 24 | 20 | 4 | 12 | 11 | 15 | 1.00E+00 | 0.80 | Retrotransposon |
| **GYPSY2** | 29 | 33 | 13 | 23 | 36 | 38 | 9.97E-01 | 1.26 | Retrotransposon |
| **GYPSY4** | 52 | 91 | 33 | 35 | 68 | 81 | 9.97E-01 | 1.05 | Retrotransposon |
| **GYPSY5** | 22 | 33 | 10 | 10 | 38 | 21 | 1.00E+00 | 1.05 | Retrotransposon |
| **GYPSY6** | 25 | 7 | 13 | 17 | 18 | 13 | 1.00E+00 | 1.04 | Retrotransposon |
| **GYPSY8** | 53 | 29 | 17 | 25 | 54 | 70 | 9.97E-01 | 1.50 | Retrotransposon |
| **GYPSY9** | 59 | 56 | 26 | 33 | 52 | 43 | 9.97E-01 | 0.91 | Retrotransposon |
| **HeT-A** | 1110 | 1252 | 658 | 729 | 1253 | 1494 | 9.97E-01 | 1.15 | Retrotransposon |
| **I-element** | 270 | 552 | 105 | 85 | 349 | 509 | 9.97E-01 | 1.02 | Retrotransposon |
| **Idefix** | 141 | 250 | 86 | 105 | 239 | 309 | 9.97E-01 | 1.36 | Retrotransposon |
| **INVADER** | 49 | 63 | 18 | 31 | 53 | 66 | 9.97E-01 | 1.15 | Retrotransposon |
| **INVADER2** | 47 | 145 | 19 | 22 | 103 | 158 | 9.97E-01 | 1.34 | Retrotransposon |
| **INVADER3** | 70 | 123 | 67 | 140 | 127 | 128 | 9.23E-01 | 1.51 | Retrotransposon |
| **INVADER4** | 36 | 29 | 10 | 35 | 47 | 73 | 8.52E-01 | 2.07 | Retrotransposon |
| **IVK** | 95 | 167 | 16 | 18 | 156 | 218 | 9.97E-01 | 1.40 | Retrotransposon |
| **Jockey** | 644 | 705 | 330 | 316 | 556 | 769 | 9.97E-01 | 0.98 | Retrotransposon |
| **Jockey_1** | 282 | 386 | 212 | 263 | 334 | 405 | 9.97E-01 | 1.14 | Retrotransposon |
| **JOCKEY2** | 219 | 196 | 108 | 201 | 199 | 166 | 9.97E-01 | 1.08 | Retrotransposon |
| **JUAN** | 87 | 82 | 23 | 17 | 73 | 77 | 9.97E-01 | 0.87 | Retrotransposon |
| **Max-element** | 2543 | 2091 | 1407 | 2719 | 2744 | 3012 | 9.96E-01 | 1.40 | Retrotransposon |
| **MDG3** | 288 | 418 | 104 | 143 | 437 | 505 | 9.97E-01 | 1.34 | Retrotransposon |
| **MGD1** | 298 | 346 | 134 | 214 | 382 | 580 | 9.97E-01 | 1.51 | Retrotransposon |
| **Microcopia** | 19 | 19 | 5 | 11 | 25 | 30 | 1.00E+00 | 1.52 | Retrotransposon |
| **ninja** | 43 | 39 | 26 | 44 | 52 | 67 | 9.97E-01 | 1.52 | Retrotransposon |
| **OPUS** | 1991 | 2046 | 588 | 1730 | 3516 | 4090 | 8.73E-01 | 2.02 | Retrotransposon |
| **QBERT** | 46 | 51 | 19 | 43 | 73 | 82 | 9.97E-01 | 1.68 | Retrotransposon |
| **QUASIMODO** | 47 | 53 | 17 | 17 | 39 | 62 | 9.97E-01 | 1.00 | Retrotransposon |
| **R1-2** | 21 | 23 | 11 | 12 | 11 | 17 | 1.00E+00 | 0.73 | Retrotransposon |
| **R1A1-element** | 7 | 27 | 11 | 28 | 26 | 25 | 1.00E+00 | 1.73 | Retrotransposon |
| **ROOA_LTR** | 382 | 444 | 69 | 144 | 454 | 1048 | 9.97E-01 | 1.84 | Retrotransposon |
| **ROVER** | 64 | 133 | 106 | 127 | 289 | 348 | 1.44E-01 | 2.52 | Retrotransposon |
| **Rt1a** | 32 | 54 | 50 | 11 | 43 | 36 | 9.97E-01 | 0.67 | Retrotransposon |
| **RT1B** | 179 | 193 | 47 | 110 | 227 | 167 | 9.97E-01 | 1.20 | Retrotransposon |
| **RT1C** | 24 | 27 | 7 | 17 | 27 | 20 | 1.00E+00 | 1.07 | Retrotransposon |
| **S-element** | 102 | 134 | 37 | 69 | 97 | 136 | 9.97E-01 | 1.10 | Retrotransposon |
| **S2** | 21 | 29 | 17 | 25 | 23 | 33 | 1.00E+00 | 1.20 | Retrotransposon |
| **SPRINGER** | 723 | 876 | 409 | 655 | 1035 | 1186 | 9.97E-01 | 1.43 | Retrotransposon |
| **STALKER2** | 450 | 745 | 348 | 130 | 178 | 214 | 1.07E-03 | 0.34 | Retrotransposon |
| **TABOR** | 149 | 240 | 54 | 71 | 169 | 258 | 9.97E-01 | 1.12 | Retrotransposon |
| **TARTC** | 106 | 189 | 51 | 34 | 104 | 141 | 9.97E-01 | 0.81 | Retrotransposon |
| **TOM1_LTR** | 72 | 75 | 91 | 110 | 93 | 89 | 9.97E-01 | 1.23 | Retrotransposon |
| **Transpac** | 569 | 833 | 296 | 273 | 953 | 973 | 9.97E-01 | 1.29 | Retrotransposon |
| **X-ELEMENT** | 67 | 115 | 21 | 15 | 58 | 102 | 9.97E-01 | 0.86 | Retrotransposon |
| **ZAM** | 24 | 41 | 29 | 7 | 34 | 28 | 9.97E-01 | 0.73 | Retrotransposon |
| **1360** | 1126 | 1263 | 543 | 916 | 1226 | 1333 | 9.97E-01 | 1.18 | DNA |
| **BARI1** | 82 | 78 | 45 | 64 | 99 | 77 | 9.97E-01 | 1.16 | DNA |
| **Hobo** | 365 | 463 | 237 | 139 | 384 | 390 | 9.97E-01 | 0.86 | DNA |
| **Pogo** | 720 | 678 | 801 | 792 | 889 | 741 | 9.97E-01 | 1.10 | DNA |
| **HB** | 161 | 215 | 40 | 53 | 167 | 173 | 9.97E-01 | 0.94 | DNA |
| **FB** | 200 | 185 | 146 | 197 | 237 | 244 | 9.97E-01 | 1.27 | DNA |
| **Hopper** | 24 | 21 | 19 | 16 | 31 | 22 | 1.00E+00 | 1.05 | DNA |
| **INE1** | 164 | 208 | 96 | 104 | 196 | 207 | 9.97E-01 | 1.08 | DNA |
| **MARINER2** | 97 | 78 | 69 | 104 | 103 | 96 | 9.97E-01 | 1.24 | DNA |
| **P-element** | 432 | 405 | 285 | 322 | 421 | 314 | 9.97E-01 | 0.94 | DNA |
| **TC1** | 126 | 156 | 29 | 43 | 90 | 147 | 9.97E-01 | 0.89 | DNA |
| **TC1-2** | 59 | 133 | 55 | 44 | 103 | 161 | 9.97E-01 | 1.24 | DNA |
| **TRANSIB2** | 123 | 102 | 49 | 71 | 115 | 117 | 9.97E-01 | 1.10 | DNA |
| **TRANSIB3** | 90 | 166 | 44 | 136 | 205 | 231 | 8.73E-01 | 1.90 | DNA |
| **TRANSIB4** | 7 | 20 | 9 | 7 | 11 | 20 | 1.00E+00 | 1.07 | DNA |

**Figure 1-Table supplement 2 (FOXO null).** FOXO null TE expression. Counts have been normalized by median ratio normalization in DESeq2. Adjusted p value calculated by DEseq2 is provided.

| TE | $\Delta 94\_6d\_1$ | $\Delta 94\_6d\_2$ | $\Delta 94\_6d\_3$ | $\Delta 94\_31d\_1$ | $\Delta 94\_31d\_2$ | $\Delta 94\_30d\_3$ | adjusted p value | Fold Change | Class |
| --- | --- | --- | --- | --- | --- | --- | --- | --- | --- |
| **17.6** | 28 | 67 | 58 | 81 | 90 | 73 | 1.50E-01 | 1.60 | Retrotransposon |
| **297** | 194 | 506 | 164 | 949 | 1081 | 270 | 1.78E-01 | 2.66 | Retrotransposon |
| **412** | 125 | 256 | 223 | 284 | 129 | 58 | 6.18E-01 | 0.78 | Retrotransposon |
| **1731** | 354 | 427 | 645 | 834 | 1605 | 2198 | 5.00E-05 | 3.25 | Retrotransposon |
| **ACCORD** | 1077 | 1259 | 1875 | 2598 | 4876 | 1519 | 2.47E-02 | 2.14 | Retrotransposon |
| **aurora-element** | 19 | 16 | 22 | 9 | 13 | 22 | 5.68E-01 | 0.74 | Retrotransposon |
| **BAGGINS** | 33 | 33 | 29 | 47 | 40 | 34 | 4.82E-01 | 1.29 | Retrotransposon |
| **Beagle** | 112 | 211 | 270 | 265 | 328 | 248 | 2.41E-01 | 1.42 | Retrotransposon |
| **Beagle2** | 8 | 21 | 25 | 29 | 53 | 22 | 1.51E-01 | 1.98 | Retrotransposon |
| **Bel** | 873 | 732 | 620 | 1128 | 894 | 1026 | 1.03E-01 | 1.37 | Retrotransposon |
| **BLOOD** | 718 | 1574 | 1252 | 1974 | 1922 | 1453 | 1.33E-01 | 1.51 | Retrotransposon |
| **BS** | 272 | 288 | 307 | 395 | 355 | 536 | 4.52E-02 | 1.48 | Retrotransposon |
| **BS3** | 47 | 46 | 54 | 41 | 38 | 41 | 5.29E-01 | 0.81 | Retrotransposon |
| **Burdock** | 888 | 1094 | 1369 | 2049 | 2463 | 1434 | 1.26E-02 | 1.78 | Retrotransposon |
| **CIRC** | 18 | 30 | 23 | 51 | 62 | 56 | 3.00E-03 | 2.43 | Retrotransposon |
| **COPIA** | 23126 | 62779 | 58084 | 126439 | 166397 | 83807 | 3.97E-03 | 2.62 | Retrotransposon |
| **CR1A** | 178 | 192 | 223 | 329 | 334 | 207 | 8.83E-02 | 1.47 | Retrotransposon |
| **Diver** | 58 | 39 | 55 | 43 | 42 | 62 | 9.25E-01 | 0.96 | Retrotransposon |
| **DIVER2** | 18 | 32 | 26 | 30 | 26 | 20 | 9.54E-01 | 1.03 | Retrotransposon |
| **DM_ROO** | 510 | 1014 | 1140 | 1350 | 1226 | 769 | 5.00E-01 | 1.26 | Retrotransposon |
| **DM88** | 38 | 54 | 43 | 44 | 42 | 20 | 5.81E-01 | 0.80 | Retrotransposon |
| **DMGYPF1A** | 166 | 141 | 165 | 274 | 300 | 277 | 4.80E-04 | 1.80 | Retrotransposon |
| **DOC** | 2636 | 1966 | 2108 | 3450 | 3628 | 2834 | 4.25E-02 | 1.48 | Retrotransposon |
| **DOC2** | 225 | 167 | 177 | 211 | 185 | 176 | 9.81E-01 | 1.00 | Retrotransposon |
| **DOC3** | 231 | 276 | 326 | 462 | 457 | 333 | 4.49E-02 | 1.51 | Retrotransposon |
| **DOC4** | 14 | 26 | 24 | 30 | 87 | 30 | 5.48E-02 | 2.32 | Retrotransposon |
| **DOC5** | 11 | 26 | 35 | 33 | 25 | 17 | 8.96E-01 | 1.08 | Retrotransposon |
| **F-element** | 385 | 382 | 421 | 529 | 446 | 361 | 5.95E-01 | 1.13 | Retrotransposon |
| **Flea** | 733 | 737 | 825 | 1431 | 1294 | 783 | 6.13E-02 | 1.53 | Retrotransposon |
| **FROGGER** | 25 | 63 | 49 | 49 | 16 | 17 | 3.25E-01 | 0.61 | Retrotransposon |
| **FW2** | 18 | 31 | 29 | 23 | 23 | 10 | 5.36E-01 | 0.74 | Retrotransposon |
| **FW3** | 15 | 30 | 30 | 21 | 13 | 10 | 2.82E-01 | 0.60 | Retrotransposon |
| **G2** | 58 | 39 | 49 | 61 | 71 | 56 | 4.40E-01 | 1.28 | Retrotransposon |
| **G3** | 13 | 17 | 28 | 12 | 7 | 9 | 1.69E-01 | 0.51 | Retrotransposon |
| **G4** | 71 | 145 | 124 | 126 | 74 | 76 | 5.83E-01 | 0.82 | Retrotransposon |
| **G5** | 98 | 73 | 93 | 97 | 88 | 101 | 8.04E-01 | 1.08 | Retrotransposon |
| **G5A** | 138 | 158 | 182 | 169 | 135 | 126 | 7.02E-01 | 0.90 | Retrotransposon |
| **G6** | 24 | 21 | 31 | 37 | 59 | 57 | 2.96E-02 | 2.02 | Retrotransposon |
| **Gate** | 166 | 121 | 166 | 362 | 484 | 472 | 1.00E-08 | 2.90 | Retrotransposon |
| **GTWIN** | 14 | 12 | 16 | 20 | 39 | 16 | 2.10E-01 | 1.78 | Retrotransposon |
| **GYPSY10** | 90 | 135 | 101 | 118 | 84 | 83 | 6.83E-01 | 0.88 | Retrotransposon |
| **GYPSY11** | 10 | 7 | 24 | 12 | 11 | 10 | 1.00E+00 | 0.81 | Retrotransposon |
| **GYPSY2** | 29 | 29 | 21 | 36 | 37 | 42 | 2.88E-01 | 1.46 | Retrotransposon |
| **GYPSY4** | 41 | 50 | 56 | 67 | 82 | 49 | 3.59E-01 | 1.35 | Retrotransposon |
| **GYPSY5** | 4 | 6 | 11 | 16 | 15 | 8 | 1.00E+00 | 1.85 | Retrotransposon |
| **GYPSY6** | 12 | 20 | 25 | 27 | 26 | 17 | 6.08E-01 | 1.28 | Retrotransposon |
| **GYPSY8** | 38 | 25 | 28 | 12 | 25 | 31 | 5.43E-01 | 0.75 | Retrotransposon |
| **GYPSY9** | 37 | 32 | 54 | 42 | 13 | 13 | 2.27E-01 | 0.56 | Retrotransposon |
| **HeT-A** | 1033 | 1077 | 1180 | 1182 | 1283 | 798 | 9.78E-01 | 0.99 | Retrotransposon |
| **I-element** | 224 | 201 | 197 | 418 | 372 | 208 | 5.26E-02 | 1.61 | Retrotransposon |
| **Idefix** | 72 | 136 | 193 | 243 | 352 | 127 | 1.08E-01 | 1.81 | Retrotransposon |
| **INVADER** | 46 | 57 | 42 | 35 | 34 | 29 | 2.34E-01 | 0.68 | Retrotransposon |
| **INVADER2** | 38 | 66 | 77 | 140 | 192 | 118 | 9.10E-04 | 2.50 | Retrotransposon |
| **INVADER3** | 150 | 176 | 185 | 241 | 336 | 196 | 7.21E-02 | 1.51 | Retrotransposon |
| **INVADER4** | 19 | 34 | 37 | 61 | 77 | 47 | 2.58E-02 | 2.07 | Retrotransposon |
| **IVK** | 53 | 55 | 45 | 125 | 225 | 224 | 1.00E-07 | 3.73 | Retrotransposon |
| **Jockey** | 848 | 596 | 653 | 831 | 1122 | 1106 | 6.34E-02 | 1.46 | Retrotransposon |
| **Jockey_1** | 579 | 539 | 644 | 519 | 542 | 409 | 3.68E-01 | 0.83 | Retrotransposon |
| **JOCKEY2** | 99 | 152 | 146 | 98 | 83 | 47 | 5.90E-02 | 0.58 | Retrotransposon |
| **JUAN** | 80 | 96 | 97 | 165 | 265 | 133 | 4.37E-03 | 2.07 | Retrotransposon |
| **Max-element** | 859 | 779 | 364 | 1190 | 807 | 781 | 3.00E-01 | 1.39 | Retrotransposon |
| **MDG3** | 712 | 708 | 553 | 925 | 753 | 408 | 8.72E-01 | 1.06 | Retrotransposon |
| **MGD1** | 178 | 350 | 425 | 553 | 833 | 473 | 1.94E-02 | 1.95 | Retrotransposon |
| **Microcopia** | 35 | 36 | 43 | 44 | 45 | 21 | 9.68E-01 | 0.98 | Retrotransposon |
| **ninja** | 33 | 28 | 39 | 17 | 33 | 23 | 4.44E-01 | 0.73 | Retrotransposon |
| **OPUS** | 1450 | 1511 | 1833 | 2025 | 2093 | 2767 | 6.66E-02 | 1.44 | Retrotransposon |
| **QBERT** | 17 | 30 | 31 | 36 | 42 | 47 | 1.74E-01 | 1.63 | Retrotransposon |
| **QUASIMODO** | 40 | 66 | 63 | 91 | 100 | 42 | 3.90E-01 | 1.39 | Retrotransposon |
| **R1-2** | 19 | 14 | 18 | 15 | 3 | 7 | 1.91E-01 | 0.50 | Retrotransposon |
| **R1A1-element** | 44 | 28 | 30 | 48 | 48 | 55 | 2.53E-01 | 1.46 | Retrotransposon |
| **ROOA_LTR** | 56 | 60 | 88 | 209 | 276 | 150 | 1.00E-05 | 3.13 | Retrotransposon |
| **ROVER** | 55 | 110 | 156 | 146 | 200 | 71 | 5.52E-01 | 1.30 | Retrotransposon |
| **Rt1a** | 27 | 41 | 32 | 55 | 54 | 30 | 3.64E-01 | 1.41 | Retrotransposon |
| **RT1B** | 136 | 171 | 167 | 183 | 212 | 143 | 6.19E-01 | 1.14 | Retrotransposon |
| **RT1C** | 26 | 19 | 25 | 19 | 19 | 20 | 6.90E-01 | 0.83 | Retrotransposon |
| **S-element** | 89 | 150 | 124 | 163 | 202 | 100 | 4.42E-01 | 1.28 | Retrotransposon |
| **S2** | 17 | 20 | 31 | 26 | 51 | 15 | 5.35E-01 | 1.38 | Retrotransposon |
| **SPRINGER** | 1363 | 1611 | 1080 | 2201 | 1600 | 1522 | 2.48E-01 | 1.31 | Retrotransposon |
| **STALKER2** | 55 | 145 | 139 | 115 | 112 | 79 | 8.19E-01 | 0.91 | Retrotransposon |
| **TABOR** | 41 | 126 | 162 | 131 | 106 | 54 | 8.24E-01 | 0.89 | Retrotransposon |
| **TARTC** | 56 | 45 | 72 | 107 | 183 | 94 | 5.13E-03 | 2.22 | Retrotransposon |
| ***TOM1_LTR*** | 28 | 52 | 76 | 66 | 105 | 69 | 2.58E-01 | 1.53 | Retrotransposon |
| ***Transpac*** | 458 | 1106 | 1447 | 2514 | 2842 | 820 | 2.94E-01 | 2.05 | Retrotransposon |
| ***X-ELEMENT*** | 68 | 45 | 55 | 62 | 53 | 55 | 9.78E-01 | 1.01 | Retrotransposon |
| ***ZAM*** | 29 | 35 | 37 | 27 | 21 | 27 | 4.38E-01 | 0.74 | Retrotransposon |
| ***1360*** | 416 | 775 | 710 | 1240 | 1224 | 734 | 4.76E-02 | 1.68 | DNA |
| ***BARI1*** | 27 | 46 | 40 | 70 | 67 | 48 | 1.12E-01 | 1.65 | DNA |
| ***Hobo*** | 90 | 185 | 138 | 175 | 188 | 155 | 4.45E-01 | 1.26 | DNA |
| ***Pogo*** | 216 | 500 | 487 | 612 | 774 | 365 | 2.69E-01 | 1.46 | DNA |
| ***HB*** | 78 | 148 | 151 | 178 | 200 | 157 | 2.08E-01 | 1.42 | DNA |
| ***FB*** | 98 | 177 | 215 | 152 | 259 | 105 | 9.03E-01 | 1.06 | DNA |
| ***Hopper*** | 9 | 37 | 20 | 30 | 41 | 23 | 4.70E-01 | 1.45 | DNA |
| ***INE1*** | 88 | 126 | 133 | 168 | 173 | 109 | 3.43E-01 | 1.31 | DNA |
| ***MARINER2*** | 84 | 113 | 114 | 98 | 81 | 89 | 6.01E-01 | 0.86 | DNA |
| ***P-element*** | 339 | 561 | 593 | 739 | 775 | 618 | 1.21E-01 | 1.43 | DNA |
| ***TC1*** | 89 | 112 | 75 | 144 | 171 | 58 | 4.30E-01 | 1.35 | DNA |
| ***TC1-2*** | 84 | 111 | 115 | 141 | 166 | 101 | 3.10E-01 | 1.32 | DNA |
| ***TRANSIB2*** | 35 | 86 | 70 | 61 | 52 | 23 | 4.49E-01 | 0.71 | DNA |
| ***TRANSIB3*** | 37 | 102 | 77 | 168 | 194 | 92 | 2.99E-02 | 2.12 | DNA |
| ***TRANSIB4*** | 14 | 14 | 12 | 23 | 17 | 9 | 7.19E-01 | 1.22 | DNA |

## Notes

### Competing Interest Statement

The authors have declared no competing interest.

